# Integrative cross-sample alignment and spatially differential gene analysis for spatial transcriptomics

**DOI:** 10.1101/2025.06.05.653933

**Authors:** Yecheng Tan, Zezhou Wang, Ai Wang, Yan Yan, Wei Lin, Qing Nie, Jifan Shi

## Abstract

Spatial transcriptomics (ST) technologies offer rich spatial context for gene expression, with varying spatial resolutions and gene coverages. However, aligning and comparing multiple ST slices, whether derived from the same or different platforms, remains challenging due to nonlinear distortions and limited spatial overlap caused by tissue processing. We present CODA, an integrative framework for **C**r**O**ss-sample alignment and spatially **D**ifferential gene **A**nalysis. CODA first learns a shared low-dimensional latent feature space across samples. Within the latent space, CODA performs global rigid alignment, applies transformer-based feature matching to identify common spatial domains, and utilizes local nonlinear refinements via large deformation diffeomorphic metric mapping, enabling a robust cross-sample comparison and extraction of spatial gene expression patterns. Benchmarking across various ST platforms demonstrates that CODA outperforms the existing methods in alignment accuracy, computational efficiency, and memory usage. Applications to vascular and brain datasets shows CODA’s ability to uncover spatially informative genes associated with disease and sleep regulation. These results highlight CODA’s broad applicability and effectiveness in ST analysis.

## Introduction

In recent years, spatial transcriptomics (ST) technologies have rapidly advanced, enabling large-scale and high-resolution profiling of gene expression while preserving spatial context within tissues^1^. Image-based methods such as seqFISH^2^, MERFISH^3^, and SMI^4^, along with sequencing-based approaches including 10X Visium^5^, Slide-seq^6^, and Stereo-seq^7^, have made it possible to capture spatial gene expression at cellular or even subcellular resolution. These technologies provide powerful tools for revealing spatial heterogeneity in tissue architecture and exploring gene functions within local microenvironments, thereby expanding the application of ST in developmental biology, cancer research, and disease mechanism studies.

Despite these advances, integrated analysis of multi-sample ST data remains challenging. First, non-overlapping sampling regions are common across experiments, making spatial correspondence difficult to establish. Second, substantial spatial shifts and nonlinear distortions introduced during tissue processing render direct coordinate-based comparison infeasible. Third, conventional analysis workflows typically identify differentially expressed genes (DEGs) based on statistical differences in expression levels^8^, rather than cross-sample variation in space. As a result, genes that maintain similar overall expression but exhibit distinct spatial distributions across different conditions may be overlooked, leading to missed spatially informative signals. Accurate alignment of spatial slices across samples therefore requires jointly capturing both molecular similarity and spatial consistency.

Several computational methods have been developed to address spatial alignment in ST data from different perspectives. PASTE^9^ employs a Gromov-Wasserstein optimal transport framework to align adjacent slices; SANTO^10^ applies a coarse-to-fine strategy using dynamic graph neural networks to identify overlapping regions; STalign^11^ and ST-GEARS^12^ leverage large deformation diffeomorphic metric mapping (LDDMM)^13^ and elastic registration to better accommodate nonlinear deformations. Among these, PASTE and SANTO primarily focus on rigid alignment and are less effective in handling complex tissue distortions. ST-GEARS, although capable of nonlinear alignment, suffers from high computational cost, limiting its scalability. More importantly, most existing methods focus solely on spatial alignment, without supporting downstream spatial analysis, thereby limiting their ability to characterize spatial expression patterns or condition-specific spatial variability across multiple ST slices. While a recent method, CAST^14^, has begun to jointly consider alignment and spatially differential gene analysis, there remains a lack of consistent and scalable approaches for analyzing spatial heterogeneity across different conditions in multi-sample ST data.

In this study, we present CODA, short for ‘‘integrative **C**r**O**ss-sample alignment and spatially **D**ifferential gene **A**nalysis for spatial transcriptomics’’. CODA addresses three key challenges—common domain identification, fast spatial alignment and spatially differential gene detection—by embedding multiple ST slices into a shared low-dimensional latent feature space. CODA first performs rigid global alignment by identifying nearest neighbors in the feature space and estimating an optimal rotation and translation using singular value decomposition. To enhance the consistency on spatial region analysis, CODA incorporates a common domain identification module. It renders the low-dimensional representations as RGB images and extracts shared spatial regions by matching visual features between source and target slices. Cells within the common domain are reselected for subsequent alignment and analysis. It then performs local alignment at the coordinate level by applying multi-channel LDDMM to construct velocity fields in the latent space, learning smooth and invertible nonlinear transformations. Based on the aligned structure, we introduce a spatial cross-correlation metric to identify spatially differential genes (SDGs) across conditions. We systematically evaluate CODA on datasets generated from multiple ST platforms and show that it outperforms existing methods in terms of alignment accuracy, computational efficiency, and memory usage. Furthermore, in applications to vascular tissue and mouse brain data, CODA identifies key genes associated with disease states and sleep regulation, demonstrating its broad utility and biological relevance in ST research.

## Results

### Overview of CODA

CODA consists of two core components: spatial alignment and spatial analysis. The alignment module establishes a unified coordinate system across samples, providing a robust foundation for subsequent spatial comparisons. Specifically, CODA decomposes the alignment task into three stages: global alignment, common domain identification, and local alignment. By jointly transforming multiple ST datasets into multi-channel feature representations, these tasks are naturally reformulated as point cloud registration, image registration, and image feature matching, respectively. Once spatial alignment is completed, CODA introduces the concept of spatial cross-correlation to enable cross-sample spatial analysis of gene expression patterns based on the aligned coordinates.

Specifically, CODA uses ST slices generated from either the same or different experimental platforms as inputs, including both sequencing-based (e.g., 10X Visium) and imaging-based (e.g., MERFISH) technologies (Fig. 1a). This cross-platform compatibility enables CODA to align and compare datasets with heterogeneous spatial resolutions. To eliminate batch effects between datasets, CODA first applies existing integration tools such as Seurat^15^ or ComBat^16^ to integrate gene expression across samples. The integrated expression matrix is then projected into a shared *d* - dimensional feature space (Fig. 1b). At this stage, each ST dataset can be interpreted as a 2D point cloud, where each point is embedded with a *d*-dimensional gene expression feature. Based on this representation, CODA performs global alignment by identifying nearest neighbors between source and target slices in the feature space. An optimal rigid transformation —including rotation and translation—is then computed using a singular value decomposition-based approach (Fig. 1c).

**Figure 1.**
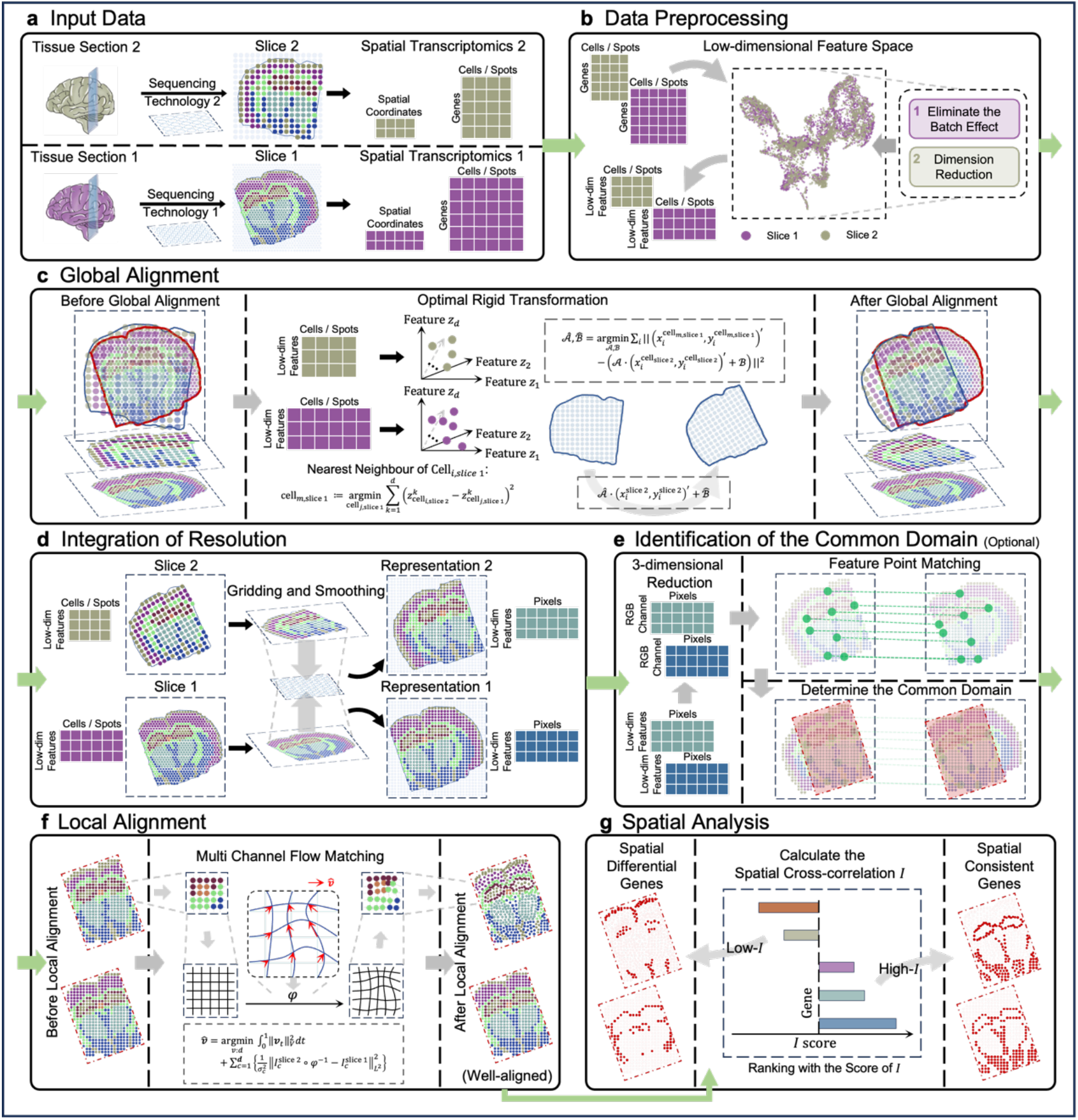
Pipeline of CODA. (a) Input data. CODA accepts two ST slices—source and target slices—from the same or different experimental platforms, which could be obtained by different sequencing technology (e.g., sequencing- or imaging-based) with diverse spatial resolutions. (b) Preprocessing. The gene expression matrices are batch-corrected and projected into a shared low-dimensional feature space. (c) Global alignment. For each cell in the source slice, its nearest neighbor in the target slice is identified in the feature space. An optimal rigid transformation—including rotation and translation—is then estimated. (d) Resolution integration. Spatial locations from both slices are embedded into a shared coordinate grid to produce multi-channel low-dimensional representations. (e) Common domain identification. A transformer-based keypoint matching model is used to identify correspondences between slices, enabling extraction of a common spatial domain. (f) Local alignment. Nonlinear geometric transformations are estimated using multi-channel large deformation diffeomorphic metric mapping (LDDMM). (g) Spatial analysis. Based on aligned spatial coordinates, CODA computes a spatial cross-correlation index to identify spatially consistent genes (SCGs) and spatially differential genes (SDGs) across conditions.

To identify a common spatial domain across slices, CODA renders the low-dimensional feature representations as multi-channel images, where each pixel encodes gene expression at a specific spatial location (Fig. 1d). When the feature dimension is 3, these representations can be interpreted as RGB images, allowing CODA to leverage image-based matching strategies. CODA adopts a transformer-based key-point matching strategy similar to LightGlue^17^, a lightweight attention model for robust correspondence detection. This enables common domain identification between slices via keypoint matching in the RGB-like representations (Fig. 1e). CODA then performs local alignment using multi-channel LDDMM on the constructed spatial representations, allowing for the alignment of nonlinear deformations and the unification of spatial scales across samples (Fig. 1f).

To allow the cross-sample analysis of gene expression more interpretable, CODA introduces a spatial cross-correlation metric that quantifies the similarity of each gene’s spatial distribution across aligned slices. Based on this metric, genes with high spatial cross-correlation are defined as spatially consistent genes (SCGs), often reflecting conserved localization patterns and known cell-type markers. In contrast, genes with low or negative spatial cross-correlation are defined as spatially differential genes (SDGs), capturing condition-specific spatial shifts that may reflect underlying biological variation. This framework enables the systematic identification of genes whose spatial organization is preserved or altered between conditions, facilitating downstream biological interpretation (Fig. 1g; see Methods for details).

### Benchmark

To evaluate the performance of CODA, we conducted a comparative benchmark against four existing methods—PASTE, STalign, ST-GEARS, and SANTO—on the task of global spatial alignment, which involves only rigid transformations (i.e., translation and rotation). Although CODA supports nonlinear transformations (local alignment), it is restricted to rigid alignment in this benchmark to ensure a fair comparison. Notably, STalign was evaluated using its default setting, which includes nonlinear deformation modeling, as its software does not support a purely rigid mode. The benchmark was conducted on three representative ST datasets generated by 10X Visium, MERFISH, and Stereo-seq, each differing in spatial resolution and sequencing depth. These datasets provide a diverse and realistic evaluation across different platforms. For each dataset, we selected two slices that are likely adjacent in the biological tissue, simulating the common task of pairwise slice alignment (see the original slices in Supplementary Fig. 3.1). Methods were compared in the aspects of alignment accuracy, computational efficiency, and memory usage (Supplementary Tables 2.1, 2.2, and 2.3, see preprocessing details in Supplementary Note 1.1).

#### CODA aligns ST slices with spot-level resolution

First, we compared the performance of different methods on the 10X Visium human dorsolateral prefrontal cortex (DLPFC) dataset^18^, a sequence-based technology with spot-level resolution. We selected two pairs of slices (ID151673 and ID151676) from an adult donor for experimental analysis. After normalizing the coordinates to a range of -1 to 1, these two slices were aligned in their initial state. To simulate rotational deviation, we manually rotated ID151673 by 180 degrees around the origin. To quantify these results, we calculated the expression correlation score (ECS), the spatial consistency index (SCI) and a rotation recovery score (RRS) designed based on the error in rotational angle (ranging from 0 to 1, where higher values indicate greater accuracy). Before alignment, the baseline performance across all metrics was low, with the ECS, SCI, and RRS for the two slices measured at 0.30, 0.18, and 0, respectively (Supplementary Table 2.1).

Among all methods, STalign failed to improve alignment quality, likely due to its reliance on single-cell resolution data for estimating cell density fields (Fig. 2a). Its performance remained consistently low across all metrics (Fig. 2d and Supplementary Table 2.1). PASTE and SANTO partially corrected the misalignment (Fig. 2a), but both remained deviated from the true orientation. They achieved moderate expression correlation scores (ECS of 0.32 and 0.34), spatial consistency indices (SCI of 0.35 and 0.37), and suboptimal rotation recovery scores (RRS of 0.73 and 0.76). These methods might be affected by residual redundancy in the gene expression space, as selection of the top 1,000 highly variable genes cannot fully eliminate noise. In contrast, CODA and ST-GEARS successfully recovered the original orientation, producing the most accurate alignments. Both achieved the highest ECS (0.37), high SCI values (0.84 and 0.86), and near-perfect RRS (0.98 and 0.99), indicating successful alignment even under extreme rotation (Fig. 2a).

**Figure 2.**
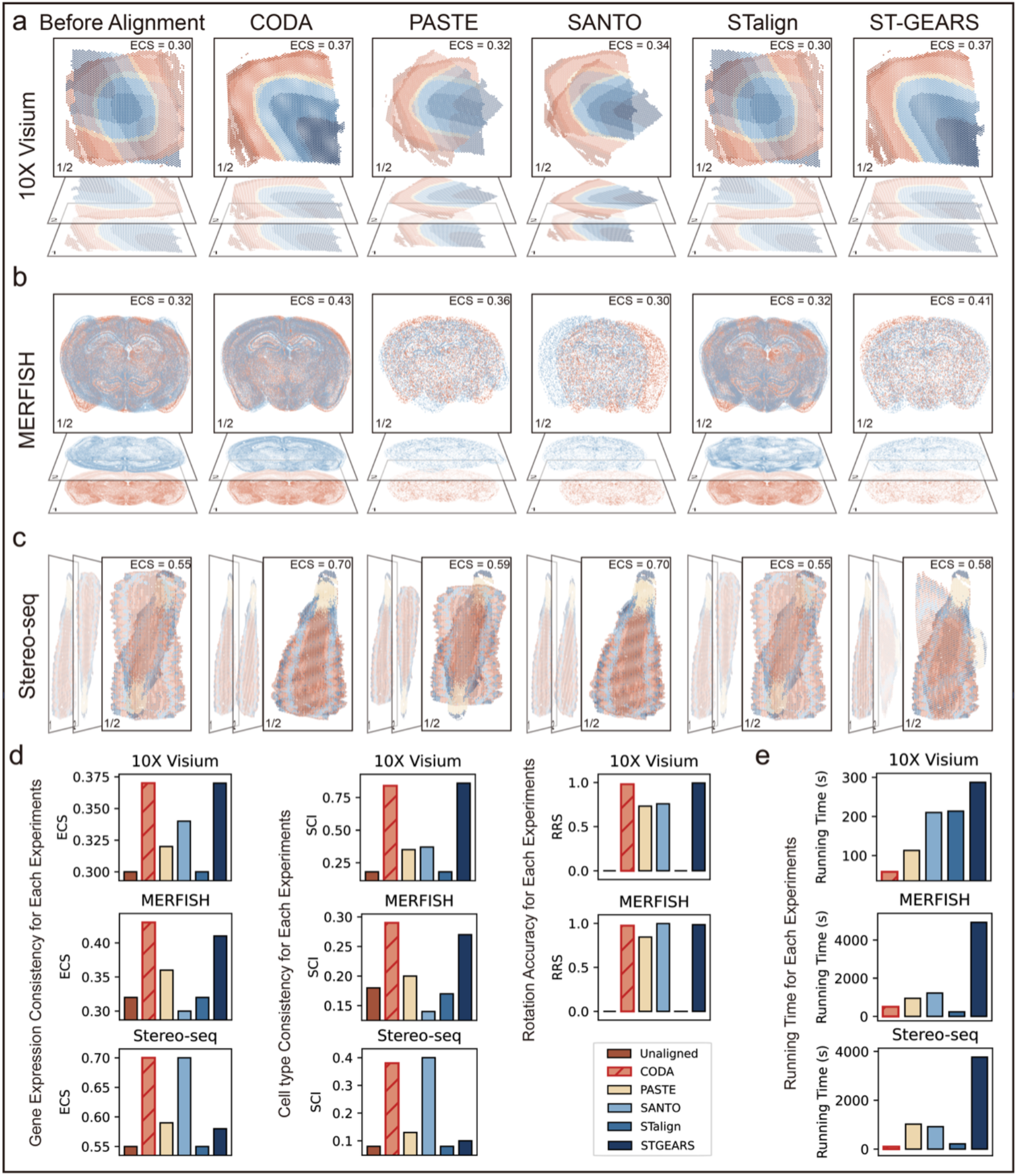
Global alignment benchmarking of CODA, PASTE, STalign, SANTO, and ST-GEARS on various ST datasets with different spatial resolutions. (a) Alignment results on the Visium DLPFC dataset. The two slices differ by 180 degrees prior to alignment. CODA and ST-GEARS achieved the highest ECS (0.37), while PASTE (0.34) and SANTO (0.32) achieved partial alignment with residual angular deviations. STalign, not designed for spot-level data, failed to align the slices. Colors represent cortical layers. Throughout all experiments, slice 2 consistently served as the source slice and slice 1 as the target. (b) Alignment results on the MERFISH mouse brain dataset. The two slices also differ by 180 degrees. Due to memory constraints, PASTE, SANTO, and ST-GEARS aligned on 10,000-cell subsets, while CODA and STalign processed the full 100,000 cells. CODA and ST-GEARS achieved the highest ECS (0.43 and 0.41, respectively); PASTE exhibited slight angular error (ECS = 0.36), SANTO introduced a translation shift, and STalign performed poorly without landmarks. Red and blue indicate cell positions from slice 1 and slice 2. (c) Alignment results on the Stereo-seq maize dataset. CODA and SANTO achieved the highest ECS (0.70), while PASTE and STalign made minimal adjustments. ST-GEARS incorrectly applied a 90-degree rotation. Colors represent different cell types. (d) Accuracy metrics (ECS, SCI, RRS) across the three benchmarks. (e) Runtime comparison for all methods of different dataset.

#### CODA aligns single-cell-resolution ST slices with low memory-cost

We next evaluated CODA on a large-scale MERFISH dataset of the mouse brain^19^, an image-based ST technology with single-cell resolution. Two spatially adjacent slices (C57BL6J-638850.38 and C57BL6J-638850.40; hereafter referred to as slice 38 and slice 40) were selected for alignment. Coordinates were normalized to [-1,1], and the slices were initially assumed to be aligned. The ECS and SCI scores of the untransformed slices were 0.44 and 0.31, which defined the upper bound of alignment performance for this pair under rigid transformation. To test the sensitivity of alignment, we applied a small 5-degree rotation to slice 38 and found that ECS and SCI dropped to 0.30 and 0.33, indicating that even minor angular deviations can significantly degrade alignment metrics. Next we applied a full 180-degree rotation to slice 38 (with the ECS and SCI dropped to 0.32 and 0.18) and evaluated alignment performance across methods. Given the large number of cells (>100,000 per slice), PASTE, SANTO, and ST-GEARS exceeded memory limits (requiring 57GB, 403GB, and 116GB, respectively). To ensure comparability, we uniformly subsampled 10,000 cells per slice for all methods, with negligible differences observed before and after sampling.

Surprisingly, STalign, despite being designed for single-cell resolution data, failed to correct the rotation and instead applied local nonlinear warping (Fig. 2b). This led to no improvement in expression correlation (ECS = 0.32) and even a decline in spatial consistency (SCI = 0.17), with a rotation recovery score (RRS) of 0 (Supplementary Table 2.2). These results suggest that STalign might strongly depend on external landmarks, which were not provided in our setup. PASTE partially corrected the angular deviation (Fig. 2b), yielding an RRS of 0.84. However, it exhibited a residual 30-degree misalignment and lower expression-based scores (ECS = 0.36, SCI = 0.20). Both SANTO and ST-GEARS successfully recovered the intended orientation, achieving high RRS values (0.99 and 0.98, respectively). Nonetheless, SANTO introduced a small translational shift, resulting in lower ECS (0.30) and SCI (0.14), suggesting that although it precisely corrected the rotation, it lacked accuracy in overall spatial positioning. ST-GEARS produced results close to CODA (ECS = 0.41, SCI = 0.27), but additionally performed an unnecessary horizontal flip, likely due to its 3D registration assumptions. CODA, in contrast, corrected the rotation without introducing additional transformations (Fig. 2b), yielding the highest ECS (0.43) and SCI (0.29) among all methods (Supplementary Table 2.2).

#### CODA robustly aligns high-resolution Stereo-seq maize slices across species and platforms

We evaluated CODA on a subcellular-resolution maize dataset^20^ generated by Stereo-seq to further assess performance across species and spatial resolutions. Two biological replicates (rep1 and rep2) were selected for alignment. Unlike previous benchmarks, these slices exhibited natural, unannotated spatial misalignment, making it infeasible to compute RRS. To estimate a performance ceiling, we applied a small 5-degree rotation to rep1 and recorded baseline scores of ECS = 0.72 and SCI = 0.41.

Among the tested methods, ST-GEARS applied an incorrect ∼90-degree rotation, indicating difficulty in rotational calibration on high-resolution, plant-derived samples (Fig. 2c). PASTE and STalign made only minimal adjustments, resulting in poor alignment quality. In contrast, SANTO and CODA both achieved high expression-based scores, with ECS = 0.70 and SCI values of 0.38 and 0.40, respectively, indicating strong alignment performance under rigid transformations without prior annotation (Fig. 2c and Supplementary Table 2.3).

#### CODA achieves high time-efficiency for large-scale datasets

We compared the time consumption of CODA with conventional methods, especially for large-scale datasets (details in Supplementary Tables 2.1, 2.2 and 2.3 as well as in the “Implementation Environment” section of the Methods). On the uniformly subsampled 10,000 cells of the MERFISH dataset, CODA completed the alignment in 12.4 seconds, with a peak CPU memory increase of only 113 MB, substantially outperforming PASTE (950s), SANTO (1,225s), and ST-GEARS (4,930s). Notably, even on the full MERFISH dataset without subsampling, CODA completed the alignment in just 510 seconds— surpassing all competing methods in speed, despite their evaluation being limited to the smaller subset. Notably, PASTE was GPU-accelerated, while SANTO and ST-GEARS ran on CPU. On the same computing setup, PASTE emerged as the most time-consuming method. While STalign was good in both runtime and memory usage, its lack of rotational correction and relatively poor alignment accuracy further highlighted the advantage of CODA’s image-based design, which balances performance, speed, and scalability.

### CODA scales to robust global and local alignment across platforms and multiple spatial slices

To evaluate CODA’s capacity for comprehensive spatial alignment, including both global and local alignment, we applied it to ST slices generated using both consistent and heterogeneous technologies. Specifically, we tested five experimental conditions: (1) two mouse hemisphere slices from 10X Visium^21^, (2) two from STARmap^22^, (3) two from RIBOmap^23^, (4) a cross-platform comparison between 10X Visium^21^ and MERFISH^19^, and (5) a multi-slice alignment task involving five Visium slices from the same dataset as (1). These datasets differ in spatial resolution, throughput, and molecular modalities, providing a comprehensive benchmark for assessing CODA’s robustness and generalizability (Supplementary Fig. 3.1).

We first examined alignment within the same technology. In the 10X Visium slices, which displayed a strong angular discrepancy, CODA applied global rotation correction followed by nonlinear refinement (Fig. 3a). The SCI increased from 0.30 (unaligned) to 0.44 after global alignment, and further to 0.53 after local alignment.

**Figure 3.**
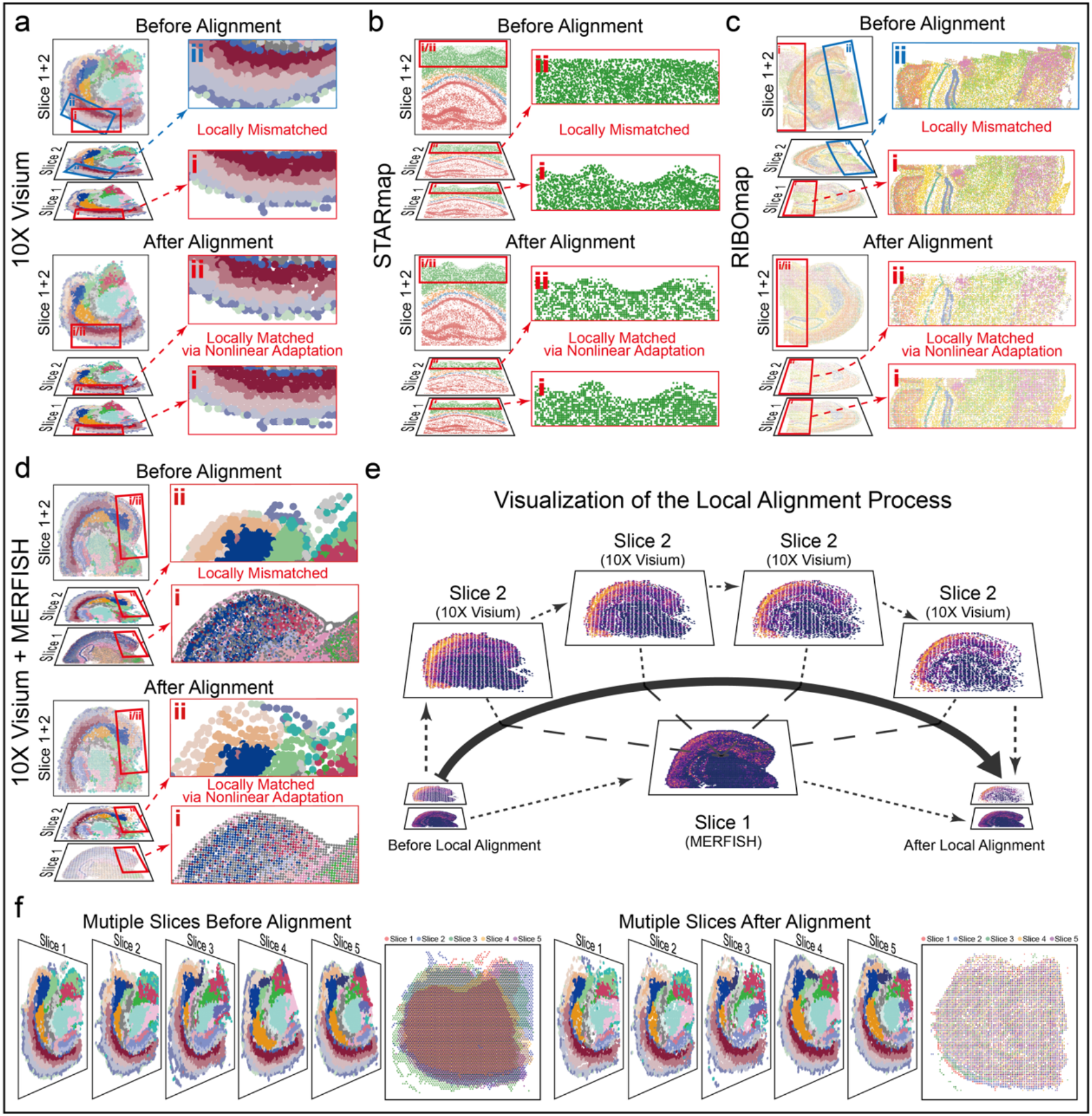
Local alignment of ST slices across technologies using CODA. (a) Alignment of two 10X Visium slices. Prior to alignment, the slices exhibited substantial angular and structural differences. CODA corrected the rotation and refined local deformations, resulting in improved spatial consistency. Colors represent different cell types. (b) Alignment of two STARmap slices. Despite minimal initial misalignment, local registration slightly improved spatial consistency across brain regions (e.g., Alveus, Corpus Callosum, Cortex). Colors denote annotated regions. (c) Alignment of two RIBOmap slices. The slices displayed significant rotational and shape discrepancies. CODA resolved both via global and local alignment, substantially improving spatial overlap. (d) Cross-technology alignment between 10X Visium and MERFISH slices. These datasets differ in resolution, cell count, and spatial sampling. CODA progressively aligned the Visium slice to match MERFISH in global shape, enabling downstream spatial comparison. (e) Evolution of spatial alignment for the gene Lamp5. Using the Visium slice as reference, cells in MERFISH were iteratively shifted based on expression similarity, yielding shape convergence. (f) Spatial alignment of five Visium slices. Prior to alignment, the slices exhibited substantial spatial misalignment. After applying CODA, the slices were brought into consistent spatial registration.

In the STARmap dataset, which provides single-cell-resolution spatial transcriptomic data, the two slices exhibited minimal angular discrepancy, resulting in negligible changes after global alignment. However, following local alignment, the spatial consistency index (SCI) improved from 0.32 to 0.43, demonstrating CODA’s capacity to refine fine-scale spatial correspondence (Fig. 3b, Supplementary Fig. 3.2).

In contrast, the RIBOmap dataset exhibited severe rotational distortion. The initial SCI was just 0.02, which increased to 0.19 after rotation correction and to 0.24 after nonlinear alignment (Fig. 3c), confirming CODA’s ability to resolve large deformations in high-resolution spatial data.

Cross-technology alignment presents greater challenges due to differences in resolution and expression modality. To test CODA’s robustness under such conditions, we aligned mouse hemisphere slices from 10X Visium and MERFISH (Fig. 3d). Using the 490 shared genes for co-embedding, CODA progressively aligned the spatial structure of the Visium slice to match that of MERFISH. Although direct region-to-region matching was infeasible due to inconsistent annotations—especially the widespread presence of OPC-Oligo cells in MERFISH—we observed that post-alignment shapes were visually concordant. To validate the effect of refined alignment at the expression level, we computed Moran’s I scores for each gene in the Visium dataset and visualized the top-ranked spatially variable genes before and after alignment. The spatial expression patterns became more consistent across datasets after applying CODA, underscoring its ability to improve both structural and molecular correspondence (Supplementary Fig. 3.3).

To assess CODA’s scalability and robustness in multi-slice alignment, we applied it to five consecutive slices from the Visium mouse brain dataset, sourced from the same sample as the one-to-one alignment in the first experiment. These slices exhibited moderate inter-slice deformation and rotational variance, providing a realistic setting for multi-slice spatial reconstruction. CODA jointly aligned all five slices by sequentially applying global and local transformations in a reference-consistent manner (Fig. 3f). Using the first slice (ST8059048) as the reference, SCI with the remaining four slices increased from 0.39, 0.29, 0.30, and 0.14 (unaligned) to 0.51, 0.41, 0.41, and 0.34 (after alignment), respectively.

### CODA identifies spatially consistent genes (SCG) and spatially differential genes (SDG) across different ST slices

Traditional single-cell analysis often overlooks genes that exhibit distinct spatial patterns despite comparable global expression levels (Fig. 4a). To address this, we introduced a spatial cross-correlation metric to quantify the similarity of gene expression patterns across ST samples. We first applied CODA to a 10X Visium human artery dataset^24^ (Fig. 4b), comprising slices from normal and atherosclerotic tissues. After alignment, nearest-neighbor analysis showed that smooth muscle cells in the diseased artery were most frequently matched to fibroblasts in the normal artery before alignment (430 cell pairs), but to smooth muscle cells after alignment (631 cell pairs; Fig. 4f). Using the aligned coordinates, we then identified two distinct gene sets: spatially consistent genes (SCGs) that maintained conserved localization across conditions, and spatially differential genes (SDGs) that exhibited marked spatial redistribution (Fig. 4c and d). Notably, SCGs often corresponded to canonical cell-type markers, whereas SDGs revealed spatial reorganization associated with disease progression.

**Figure 4.**
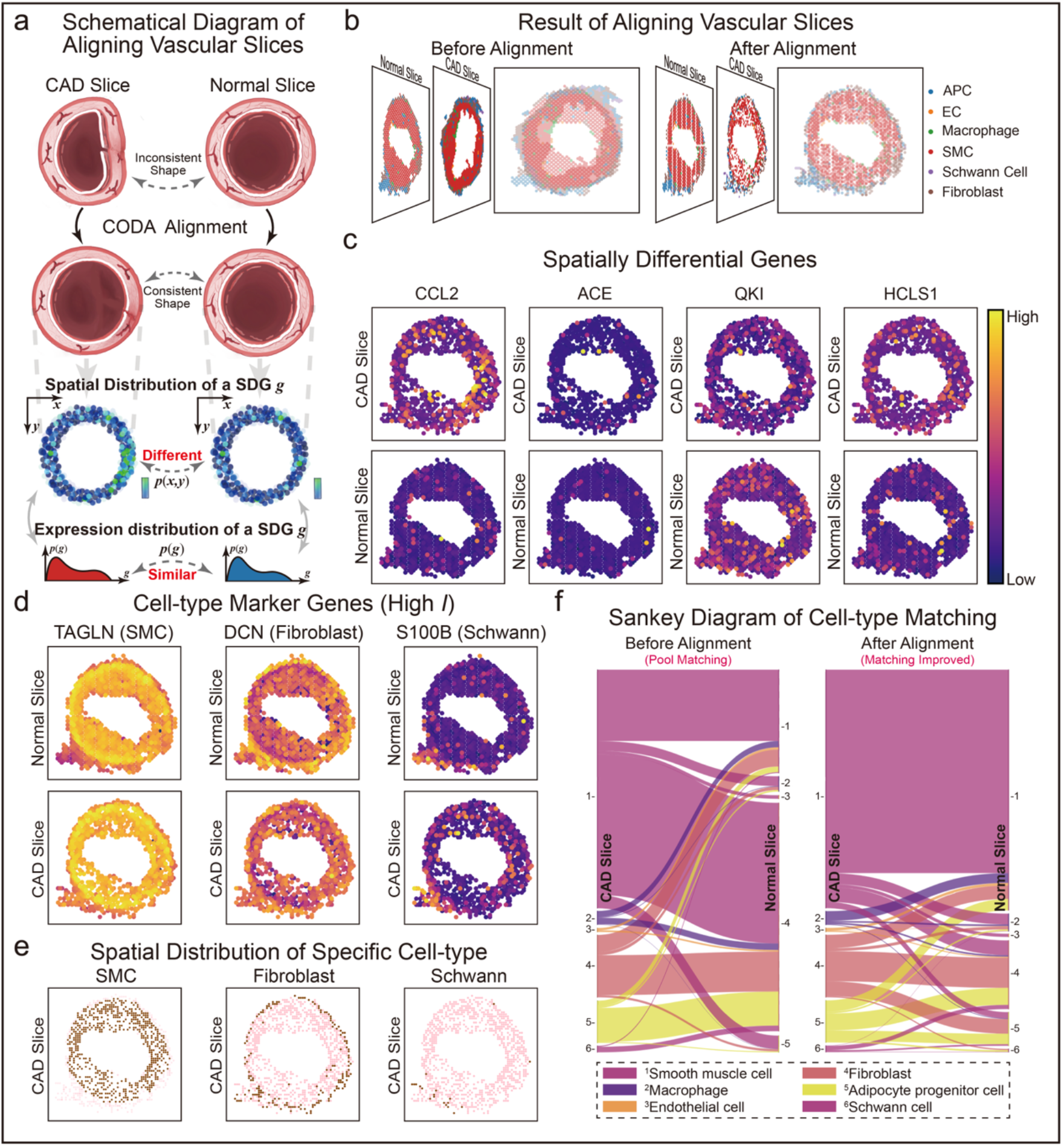
Spatial alignment and identification of spatially differential genes in arterial tissue. (a) Schematic illustration of spatially differential gene expression. The top row shows original vessel structures from a normal artery and an atherosclerotic artery, where wall damage and immune cell infiltration cause visible structural distortion in the latter. The middle row shows artificially aligned slices, emphasizing shape similarity. The bottom row highlights a spatially differential gene: although expression levels are comparable across slices, its spatial distribution differs significantly — accumulating near the damaged wall in the diseased sample but remaining diffuse in the healthy tissue. (b) Overlay of arterial slices before and after alignment. Misalignment between the atherosclerotic and normal tissues is visible prior to alignment (left), which is corrected by CODA to reveal spatial correspondence between structures (right). (c) Spatial expression of genes with low spatial cross-correlation (SDGs), reflecting condition-specific distribution patterns. Notable examples include CCL2, ACE, QKI, and HCLS1. (d) Spatial expression of genes with high spatial cross-correlation (SCGs), which reflects conserved localization patterns across slices. Examples include TAGLN (rank 2, smooth muscle cells), DCN (rank 11, fibroblasts) and S100B (rank 39, Schwann cells). (e) Spatial distributions of smooth muscle cells, fibroblasts, and Schwann cells, corresponding closely to the expression patterns of spatially consistent marker genes shown in Fig. 4d. (f) Sankey diagram showing nearest-neighbor cell-type correspondences between CAD and normal artery slices before and after alignment. Before alignment, many smooth muscle cells (SMCs) were closest to fibroblasts in the other slice. After alignment, SMCs were more consistently matched to SMCs.

To assess the biological relevance of SCGs, we examined the top-ranked genes by spatial cross-correlation index and observed that many corresponded to canonical markers of major cell types in arterial tissue (top 2%, full list in Supplementary Table 4). For example, SMC for smooth muscle cells (ranked 2), DCN for fibroblasts (ranked 11), APOD for adipocytes (ranked 4), CD36 for macrophages (ranked 36), and S100B for Schwann cells (ranked 39) all exhibited high spatial cross-correlation, indicating conserved spatial localization across samples (Fig. 4d and Supplementary Figs. 3.4 and 3.5). These genes show strong spatial clustering following alignment, as CODA adjusted the positions of similar cell types into comparable spatial neighborhoods across slices.

In contrast, SDGs—those with low spatial cross-correlation—exhibited significant differences in spatial distribution across slices (Supplementary Table 2.5). Notably, this divergence may not always be accompanied by large changes in gene expression levels, making such genes difficult to detect using traditional DEG analysis. Among the lowest-ranked genes by spatial cross-correlation (Fig. 4c, Supplementary Fig. 3.6), we identified several with known links to atherosclerosis. For instance, CCL2 (ranked 2999) is a chemokine involved in monocyte/macrophage recruitment and immune-inflammatory responses^25^; ACE (ranked 2947) is highly expressed in lipid-rich macrophages and regulates both vascular tone and lipid metabolism via PPAR α signaling^26^. Other low-ranking genes include MKI67 (ranked 3000), LY96 (ranked 2955), and CTSD (ranked 2977), all of which have been implicated in atherosclerosis-related processes^27,28,29^.

To further assess whether these genes are captured by standard expression-based methods, we performed Wilcoxon-based differential gene expression analysis, computing log fold-changes and adjusted p-values. Although ACE had the highest log fold-change (0.47, ranked 601st by absolute value), its adjusted p-value was 0.89, failing to meet statistical significance level, which is usually set at 0.05. For other SDGs, the largest significant log fold-change (CCL2) was only 0.14, well below typical thresholds used in DEG analyses. These results underscore CODA’s ability to uncover spatially distinct gene regulation patterns that are invisible to expression-level comparisons alone.

### CODA extracts common domains and identifies spatially consistent genes across slices

To further demonstrate CODA’s ability to resolve spatial variation and identify meaningful biological patterns, we applied it to mouse brain coronal slices generated by two ST platforms: CosMx Spatial Molecular Imager (SMI) ^30^ and 10x Genomics Visium^31^. The SMI dataset included two slices—one spanning a full hemisphere and the other focused on the hippocampus and cortex—while the Visium dataset contained coronal slices covering the cortex, hippocampus, and hypothalamus.

Accurate slice alignment requires sufficient overlap in both gene expression and spatial coverage. However, in practice, technical variation and experimental inconsistencies often lead to substantial differences in the detected tissue areas, particularly across platforms or batches. This issue is evident in the SMI dataset, where the two slices covered markedly different areas (17.1 mm^2^ and 33.68 mm^2^), with only partial spatial overlap (Fig. 5a). Unlike many existing methods that require fully overlapping tissue areas, CODA is able to robustly extract and align a common domain even under partial spatial overlap. By visualizing both slices as RGB-encoded spatial representations and applying a transformer-based feature matching strategy, CODA detects and aligns corresponding keypoints across slices (Fig. 5b). This process enables CODA to define a common domain between the two slices (Fig. 5b), which was further confirmed through visual inspection (Fig. 5c). Extracting this shared region led to marked improvements in downstream alignment: the ECS and SCI increased from 0.41 and 0.05 to 0.51 and 0.16, respectively.

**Figure 5.**
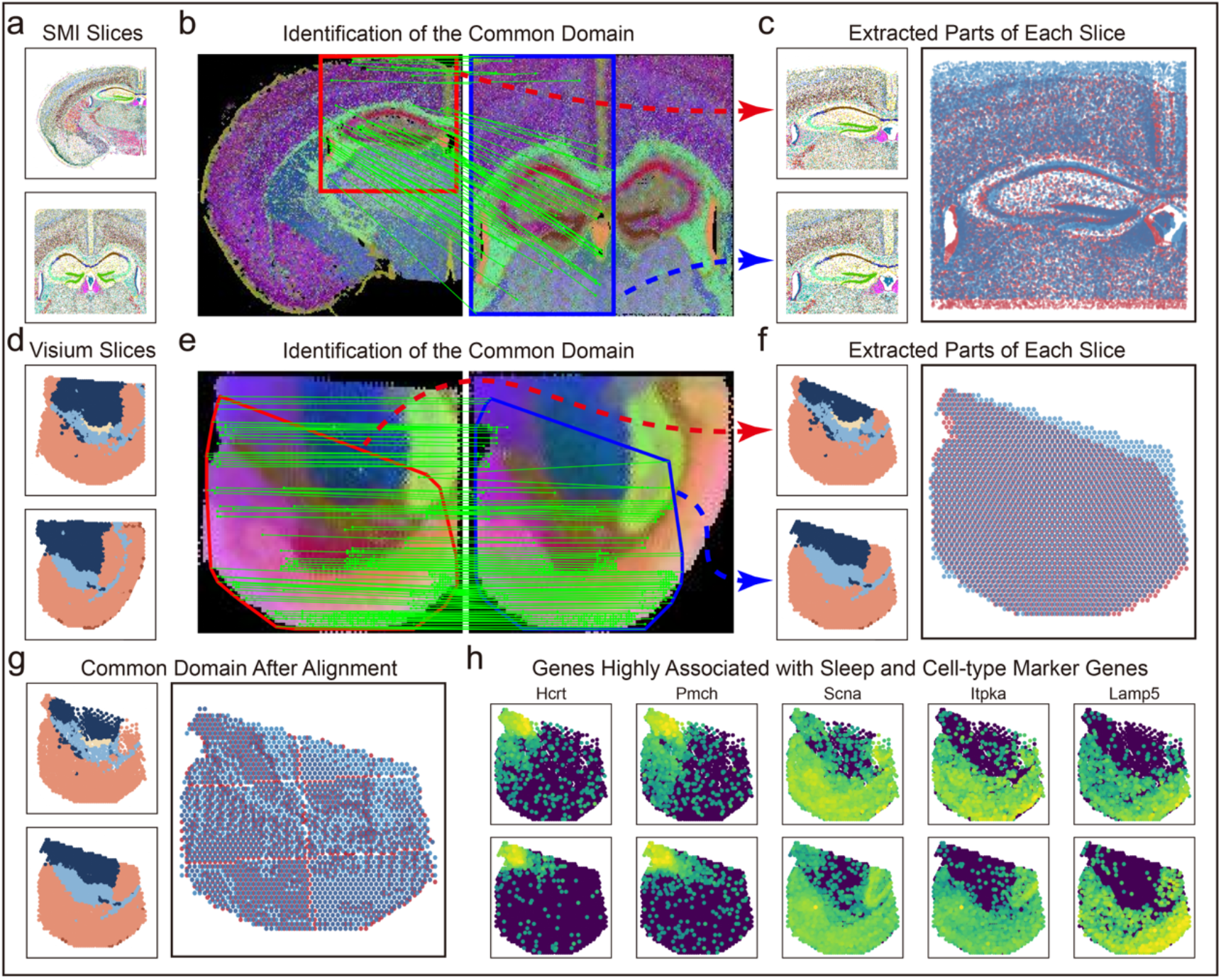
Identification of common domains and spatial analysis in mouse brain datasets. (a) Two ST slices generated using the CosMx Spatial Molecular Imager (SMI) platform. The coverage areas of the two slices partially overlap. (b) Identification of the common domain between the two SMI slices using spatial feature matching. (c) Extracted common domain from the two SMI slices. (d) Original mouse brain coronal slices from the 10X Visium dataset. (e) Identification of the common domain between the two Visium slices. (f) Extracted common domain from the Visium slices. (g) Common spatial domain identified after alignment. Compared to the unaligned slices, the extracted domains exhibit stronger overlap across samples. (h) Genes with high spatial cross-correlation index, showing consistent spatial expression patterns across slices.

To further evaluate the role of spatial cross-correlation in identifying genes with distinct spatial patterns across conditions, we applied CODA to a Visium-based mouse brain dataset designed to study the effects of Rhynchophylline (RHY) treatment. This dataset includes multiple coronal slices sampled under different doses (50 mg/kg and 100 mg/kg), time points, and biological sexes (details in SI). In the saline group at ZT4, we observed a substantial region missing from one of the slices, resulting in a spatial mismatch with the corresponding RHY-treated slice (Fig. 5d). CODA successfully extracted and aligned the common domain between the two slices (Figs. 5e–g), enabling a meaningful spatial comparison.

After alignment, we computed the spatial cross-correlation for all genes and ranked them (Supplementary Table 2.6). Genes with high spatial cross-correlation—such as *Baiap3* and *Nrgn*—exhibited consistent spatial expression across slices, and included known cell-type markers like *Lamp5* and *Itpka* (Fig. 5h). Notably, unlike atherosclerosis, RHY does not induce structural disruption in brain tissue. Consistent with this, sleep-related genes such as *Hcrt*^32^ and *Pmch*^33^ exhibited substantial changes in expression levels but maintained stable spatial localization, ranking 6th and 27th in spatial cross-correlation. In contrast, genes with low spatial cross-correlation often displayed noisy or non-specific spatial patterns despite expression differences, highlighting the importance of integrating both spatial and transcriptional signals (Supplementary Figs. 3.7 and 3.8).

### CODA enables robust alignment across diverse species and organs

To further evaluate the generalizability of CODA across different biological contexts, we applied CODA to four datasets spanning distinct species and organs.

#### CODA achieves accurate spatial alignment in plant root tissue

We first applied CODA to a ST dataset of soybean root slices^34^. CODA significantly improved inter-slice spatial consistency with the spatial consistency index (SCI) increased from 0.45 to 0.53, indicating enhanced alignment of gene expression structures across slices.

#### CODA aligns mammalian organ slices across developmental variation

We next applied CODA to gestational uterine tissues undergoing placentation^35^ and mouse embryonic tissues^36^, which exhibit complex organogenesis and layered tissue structure. In the uterine dataset, CODA corrected both global and local spatial discrepancies, resulting in an increase in SCI from 0.18 to 0.56. In the embryonic dataset, the two slices initially showed notable mismatches in tissue-to-tissue correspondence, as evident from the disorganized off-diagonal patterns in the nearest-neighbor heatmap (Fig. 6d, before alignment). After alignment, the slices reached a high degree of morphological concordance (Fig. 6c), with clear one-to-one correspondence across major anatomical regions (Fig. 6d, post-alignment), and the SCI improved substantially from 0.20 to 0.68.

**Figure 6.**
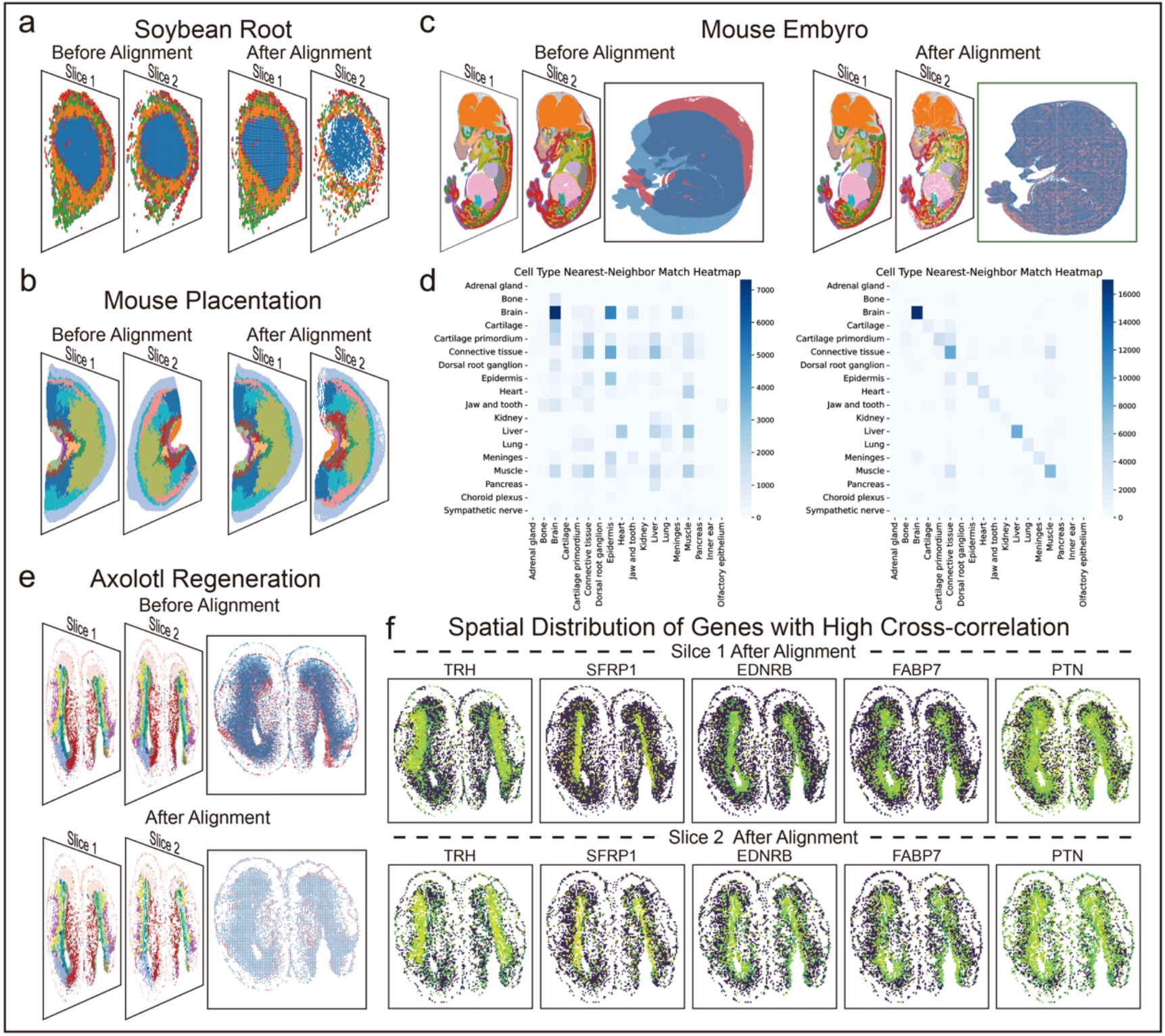
CODA generalizes across diverse biological systems and species. (a) Spatial alignment of soybean root slices before and after applying CODA. (b) Alignment of gestational uterine tissues undergoing placentation before and after CODA alignment. (c) Alignment of mouse embryonic slices before and after applying CODA. (d) Heatmaps showing nearest-neighbor cell-type correspondences between two embryonic slices before and after alignment. Prior to alignment, mismatches across anatomical regions resulted in off-diagonal patterns. After alignment, the heatmap approaches a diagonal structure, indicating improved anatomical correspondence. (e) Alignment of injured axolotl telencephalon slices before and after applying CODA. (f) Spatial expression patterns of top-ranked genes by spatial cross-correlation. These genes include known tissue-specific markers and genes previously implicated in regeneration.

#### CODA reveals spatially conserved regenerative signals in injured axolotl

We further applied CODA to regenerating axolotl limb tissue from injured axolotl telencephalon at 2 days post injury (2 DPI)^37^. CODA successfully aligned slices from the injured telencephalon, improving SCI from 0.36 to 0.46, and enabling robust identification of spatially consistent genes across sections. Among the top-ranked spatially consistent genes were *GFAP, FABP7*, and *VIM*, well-established markers of radial glia and reactive astrocytes, as well as *GAD1* and *GAD2*, which label inhibitory interneurons such as *sstIN* and *scgnIN* in the annotated cell types (Fig. 6f, Supplementary Figs. 3.9, 3.10 and Supplementary Table 2.7). In addition, we observed high spatial concordance for *TRH, EDNRB, SFRP1, RGCC*, and *PTN* — genes previously implicated in neuroendocrine signaling, neural crest–derived cell lineages, and injury response pathways (Fig. 6f, Supplementary Figs. 3.9, 3.10 and Supplementary Table 2.7). These findings suggest that CODA is capable of capturing conserved spatial programs associated with both cell-type-specific architecture and regenerative molecular cues, even in highly dynamic and heterogeneous post-injury contexts.

## Discussion

In this study, we propose CODA, a scalable and computationally efficient framework for spatial alignment and cross-sample analysis in spatial transcriptomics. CODA addresses several unaddressed challenges. Both global and local nonlinear alignment are introduced with minimal memory usage and runtime, enabling robust integration of spatial slices even in large-scale or high-resolution scenarios. CODA is able to identify shared spatial domains between samples, allowing simultaneous alignment and downstream analysis for biologically comparable regions. In addition, CODA introduces a spatial cross-correlation metric, enabling the detection of condition-specific spatial expression patterns beyond traditional expression-level comparisons.

Compared to existing alignment tools such as PASTE, STalign, SANTO, and ST-GEARS—often limited to rigid registration or are computationally intensive—CODA achieves superior alignment accuracy while maintaining high efficiency and low resource consumption. More importantly, CODA extends beyond spatial alignment by supporting downstream spatial analysis, offering a comprehensive framework for the integration and interpretation of ST datasets across diverse experimental conditions. Benchmark evaluations across multiple ST platforms, including 10X Visium, MERFISH, Stereo-seq, and STARmap, demonstrate CODA’s versatility and consistent performance across varying resolutions and data modalities.

Despite its strengths, CODA has several limitations. First, it relies on a low-dimensional latent space for alignment, which may be sensitive to feature selection and gene integration strategies. Second, the current framework adopts a pairwise or reference-based alignment paradigm; extending CODA to fully symmetric groupwise alignment remains an open direction. Third, while CODA supports spatial differential analysis, its integration with other layers of spatial omics—such as spatial proteomics or chromatin accessibility profiling (e.g., spatial ATAC-seq)—has yet to be explored. Recent advances in multimodal spatial data integration^38,39,40^ offer promising directions for extending CODA’s analytical capabilities across molecular layers.

Moreover, an increasing number of computational frameworks have been developed recently for inferring cell-cell communication (CCC) in spatial transcriptomics^41,42,43^. Integrating CODA with such CCC inference methods will enable deeper insights into spatial signaling dynamics, tissue organization, and pathology. In addition, combining CODA with gene regulatory network inference or spatial epigenomic profiling may allow for a more comprehensive understanding of spatial gene regulation. Integrating CODA with generative models will allow an estimate of missing intermediate tissue sections, and fully unsupervised groupwise alignment and automatic anchor selection will improve its autonomy and robustness. In summary, CODA serves as a generalizable and extensible framework for spatial data integration and analysis, helping to unlock the full potential of spatial omics in complex biological systems.

## METHODS

### Alignment details

#### Notations

We denote a ST slice as (***X, Y***), where ***X*** ∈ *R*^*G*×*N*^ represents the gene expression matrix of cells or spots, and ***Y*** ∈ *R*^*D*×*N*^ represents the spatial coordinate matrix. The constants *G, N* and *D* represent the number of genes, the number of cells/spots and the dimensionality of spatial coordinates (usually *D* = 2 or 3 in physical space), respectively. In the alignment task, we have a source slice (***X***_*S*_, ***Y***_*S*_) and a target slice (***X***_*T*_, ***Y***_*T*_), where 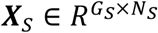 has *G*_*S*_ genes and *N*_*S*_ cells/spots while 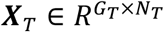 has *G*_*T*_ genes and *N*_*T*_ cells/spots. When multiple slices require alignment, we typically designate one slice as the target and sequentially align the remaining slices to this target slice in a pairwise manner.

#### Data preprocessing

We first select the common genes between the source and target slices and integrate the gene expression matrices ***X***_*S*_ and ***X***_*T*_ using methods such as Seurat, Combat, or BBKNN. We recommend Seurat or Combat, which can remove batch effect and adjust gene expression levels across different datasets effectively. The modified gene expression matrices after integration are denoted as 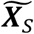 and 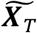 . Then, we concatenate the gene expression matrices and use UMAP to reduce the dimensionality to *d* dimensions:

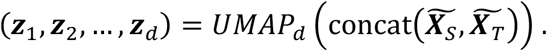

Subsequently, we normalize the *d*-dimensional data to [0,255]:

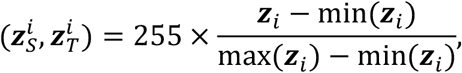

where min(***z***_*i*_) and max(***z***_*i*_) (*i* = 1,2, …, *d*) are the minimum and maximum values of the *ith* dimension, and subscripts *S* and *T* represent the *“*source slice*”* and the *“*target slice*”*, respectively. As a result, we obtain low-dimension representations for each cell/spot in the ST data.

#### Global alignment

After preprocessing, each cell/spot in source or target slice has *d* features *z*_*i*_ and two spatial coordinates *x*_*i*_ and *Y*_*i*_, which is denoted as 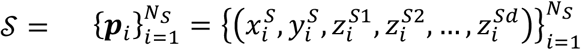 and 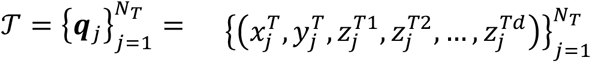, respectively. Global alignment is to find a rotational 𝒜 and a translation ℬ between source and target (rigid transformation). For each sample ***p***_*i*_ in the source slice, we find its nearest neighbor ***q***(*i*) in the feature spaces 𝒮 and 𝒯:

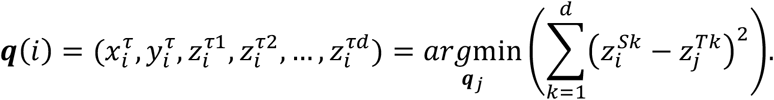

Find a rotational transformation 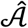 and a translational transformation 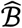 that minimize the overall distance between samples in the source slice and their nearest neighbors in the target slice in the feature space:

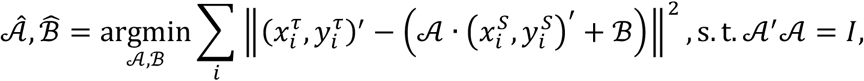

where *I* is the identity matrix and (·)^′^ denotes the matrix transpose. Center the samples in the source slice and their corresponding nearest neighbors:

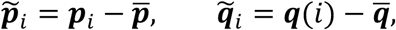

where 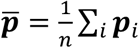and 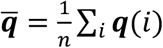 are the centroids of the source slice samples and their nearest neighbors in the target slice, respectively. Calculate their covariance matrix 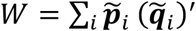, and by performing SVD decomposition *W* = *U*Σ*V*^′^, the final rotational transformation 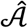 and a translational transformation 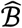 are represented by 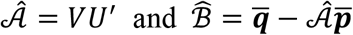, respectively.

#### Integration of Resolution

After the global alignment is done, we aim to generate the representation metrices *I*_*S*_ and *I*_*T*_ for the source and target slice in the same spatial resolution. We start by setting the spatial resolution (representation size) to *n* × *m* (default by 100 ×100, ensuring that the inserted pixel values do not overlap in the representation), and initializing *I*_*S*_ and *I*_*T*_ as *n* × *m* × *d* zero matrices. Given a set of spatial coordinates 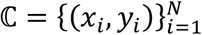 and corresponding *d* -dimensional feature values 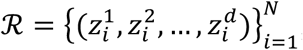, we normalize and scale the coordinates (*x*, *Y*) to the image size *n* × *m* by:

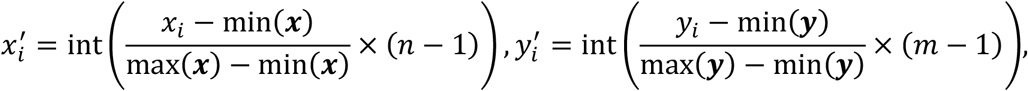

where ***x*** and ***y*** are the set of all *x*_*i*_ and *Y*_*i*_, respectively, and int(·) denotes rounding to the nearest integer. At the scaled coordinates 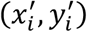, set the corresponding pixel in the presentation matrix *I* to 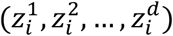. There could be many zero pixels surrounding each cell/spot in *I* resulting in visual discontinuities. Then, we perform interpolation on the representation to improve the resolution for the subsequent refined alignment. We set a neighborhood radius size *r* (default size is 1, which can be adjusted according to the specific requirements of different ST technologies used in alignment tasks), and the neighborhood *𝒩* of a pixel point *I*(*a, b*) is defined as *𝒩*>*I*(*a, b*)) = {(*i, j*), *i, j* ∈ *N*|*i* ∈ [*a* − *r, a* + *r*], *j* ∈ [*b* − *r, b* + *r*]} . We retrieve all zero pixels, and for zero pixels where the number of non-zero pixels in the neighborhood is three or more, we perform interpolation on that pixel. The interpolation is calculated as the weighted average of the non-zero pixels within the neighborhood:

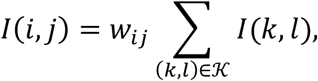

where 𝒦 ⊆ *𝒩*(*I*(*i, j*)) represents the set of all non-zero pixels within the neighborhood of pixel *I*(*i, j*), and *w*_*ij*_ are the weights corresponding to each non-zero pixel. By default, 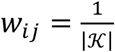, where |𝒦| is the number of elements in the set 𝒦.

#### Identification of common domains

CODA employs a similar strategy as LightGlue. First, CODA extracts feature points from the source and target representations *I*_*S*_ and *I*_*T*_, and establishes initial correspondences between them. To enhance the representation of each feature point, we employ a transformer-based architecture consisting of *L* alternating self-attention and cross attention layers. Each layer updates the feature representation *z*_*i*_ as follows:

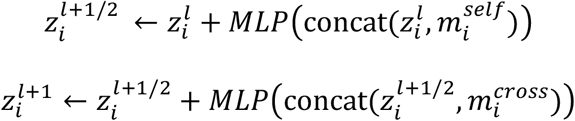

where *m*_*i*_ is the aggregated message computed from attention.

In the self-attention module, each point attends to all other points within the same presentation. The key, query and value vectors are computed as:

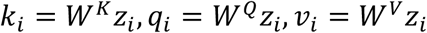

A rotation-based positional encoding *R* is added to incorporate spatial information. The attention score between points *i* and *j* is calculated as:

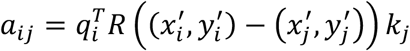

where *R* is a rotation encoding:

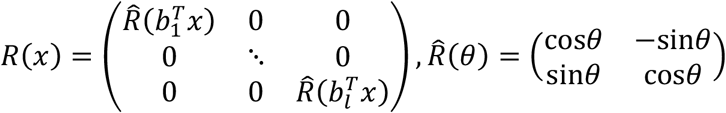

where *b*_1_, *b*_2_, …, *b*_*l*_ ∈ *R*^2^ are learnable parameters.

The output message is:

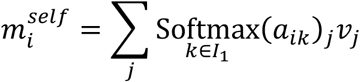

In the cross-attention module, the key and value vectors are similarly computed:

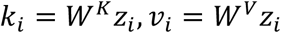

The attention score between points *i* and *j* is calculated as:

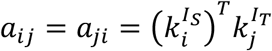

And the output message is:

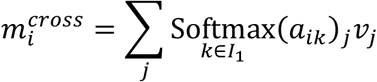

The similarity score matrix between source and target points is computed as:

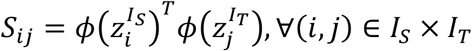

where *ϕ* is a learnable linear projection. And *S*_*ij*_ is normalized as follows:

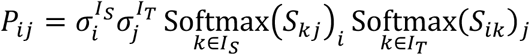

where *σ*_*i*_ = *Sigmoid*(*ϕ*(*z*_*i*_)) ∈ [0,1] . A pair of (*i, j*) is considered a valid correspondence if both points are predicted as matchable, *S*_*ij*_ is greater than a threshold *δ*, and is larger than all other entries in its row and column.

The Loss function is

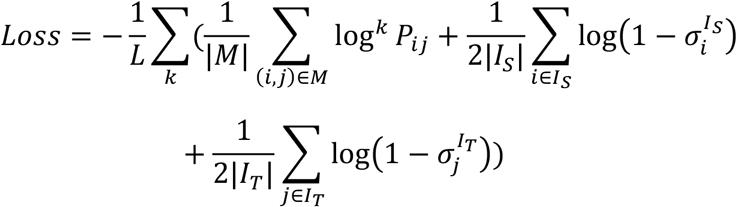

After the matched points are identified, CODA supplies two strategies to define the common domain from the set of correspondences (see Supplementary Note 1.2 for details).

#### Local alignment

After the global alignment and imaging of the slices, ensuring that both slices are within the same area, CODA performs a local alignment of the source and target slices based on nonlinear velocity fields using multi-channel LDDMM.

For two representations *I*_0_ and *I*_1_, LDDMM performs alignment by finding a diffeomorphism *φ* that satisfies 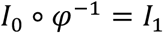, where *φ* = *ϕ*_1_ is the endpoint of the flow associated with a smooth time-dependent vector field. The evolution equation 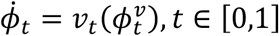 is solved for *φ*, where *v*_*t*_ is a time-varying velocity field. For given representations *I*_*S*_ = (***I***_*S*1_, ***I***_*S*2_, …, ***I***_*Sd*_) and *I*_*T*_ = (***I***_*T*1_, ***I***_*T*2_, …, ***I***_*Td*_), where ***I***_*Sc*_, ***I***_*Tc*_: Ω ⊂ ℝ^2^ → ℝ, *c* = 1,2, …, *d*, the velocity field *v*_*t*_ is determined by solving the following variational problem:

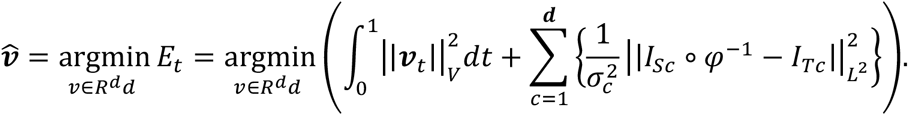

The gradient of the above loss function is:

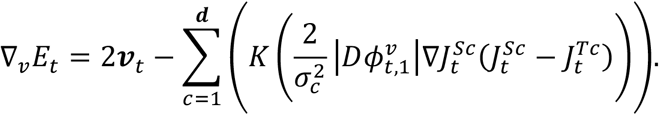

where 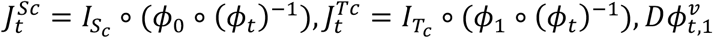 is the Jacobian of mapping 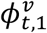 . By using the gradient descent algorithm, *v*_*t*_ can be obtained. Then, by solving the ODE 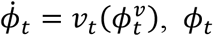 can be determined, resulting in the final *φ* = *ϕ*_1_.

After aligning *I*_*S*_ and *I*_*T*_ using multi-channel LDDMM, we determine the final positions of cells/spots in the source slice by identifying the ultimate pixel locations of their original pixels after transformation. Simultaneously, we update the positions of cells/spots in the target slice to the pixel locations corresponding to their generated presentations.

#### Identification of spatially differential genes (SDG) via spatial cross-correlation

To identify spatially differential genes (SDGs), we employed a spatial cross-correlation analysis, which modifies Moran’s *I* index to assess the correlation between aligned spatial gene expression datasets. The cross-correlation index is defined as:

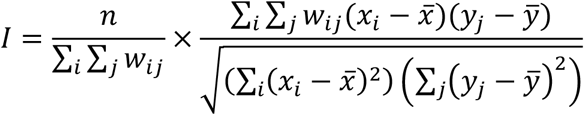

where *n* is the sample size of slice A, *x*_*i*_ and *y*_*j*_ represent the gene expression levels of gene *g* in cells *i* and *j*, respectively. 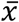 and 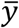 are the average expression levels of gene *g* in slices A and B, respectively. *w*_*ij*_ is the weight between cell *i* in slice A and cell *j* in slice B, defined as 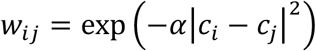, where *c*_*j*_ and *c*_*j*_ represent the aligned positions of cells *i* and *j* in their corresponding slices. With this measure, SDGs were identified as genes exhibiting low spatial cross-correlation values, indicating notable differences in spatial expression patterns across slices.

### Benchmark

#### Evaluation metrics

In the Benchmark analysis, we employed the Expression Correlation Score (ECS), Spatial Consistency Index (SCI), and Rotation Recovery Score (RRS) to evaluate the accuracy of alignment. These metrics assess different aspects of ST alignment, including expression similarity, cell type/region consistency, and spatial orientation accuracy.

### Expression Correlation Score (ECS)

The ECS measures the correlation of gene expression profiles between spatially paired points in two aligned slices. For each point *x*_*i*_ ∈ *X* in the source slice X, we identify its nearest spatial neighbor *Y*_*j*_ ∈ *Y* in the target slice Y after alignment. The ECS is calculated as the mean Pearson correlation coefficient across all paired points:

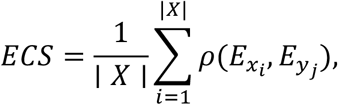

where 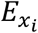 and 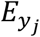are the gene expression vectors for points *x*_*i*_ and *Y*_*j*_, respectively, and *ρ* denotes the Pearson correlation coefficient. A higher ECS indicates better correspondence of gene expression.

### Spatial Consistency Index (SCI)

The SCI evaluates the consistency of cell-type/region annotations between aligned slices. For each point *x*_*i*_ ∈ *X*, we check whether its closest spatial neighbor *y*_*j*_ ∈ *Y* belongs to the same cell type. Let 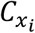 and 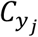 represent the cell-type annotations of *x*_*i*_ and *y*_*j*_, respectively. The SCI is defined as the proportion of matched cell types/regions:

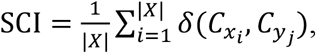

where

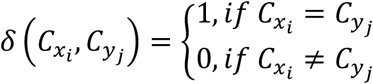

A higher SCI indicates a greater alignment of biological identities between slices.

### Rotation Recovery Score (RRS)

The RRS quantifies the accuracy of recovering the original rotation angle. Let *θ*_*pred*_ and *θ*_*true*_ denote the predicted rotation error and the ground-truth rotation error, respectively. The RRS is computed as:

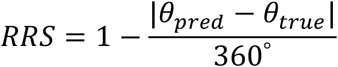

A RRS closer to 1 indicates a more accurate recovery of the applied rotation.

### Implementation Environment

All experiments were conducted on a Win11 system equipped with Intel i5-13490F and Crucial 16GB*2 DDR4-3200MHz. GPU acceleration was enabled using an NVIDIA GeForce RTX 4070 SUPER with 12 GB memory, running NVIDIA driver version 560.94.

## Data availability

The original public data used in this paper can be accessed through the following links: 1. 10X Visium dataset of the human dorsolateral prefrontal cortex (DLPFC): http://spatial.libd.org/spatialLIBD/; 2. MERFISH dataset of the mouse brain: https://portal.brain-map.org/atlases-and-data/bkp/abc-atlas; 3. Stereo-seq dataset of the developing maize ear: https://db.cngb.org/stomics/datasets/STDS0000236/summary; 4. 10X Visium dataset of the mouse brain: https://www.ebi.ac.uk/biostudies/arrayexpress/studies/E-MTAB-11114; 5. STARmap PLUS dataset of the mouse brain: https://singlecell.broadinstitute.org/single_cell/study/SCP1375; 6. RIBOmap dataset of the mouse brain: https://singlecell.broadinstitute.org/single_cell/study/SCP1835; 7. Human artery spatial transcriptomics dataset: https://doi.org/10.5281/zenodo.15132967; 8. SMI dataset of the mouse brain (CosMx platform): http://nanostring.com/CosMx-dataset; 9. Mouse brain spatial transcriptomics during sleep: https://www.ncbi.nlm.nih.gov/geo/query/acc.cgi?acc=GSE217058; 10. Soybean root nodule dataset: https://ngdc.cncb.ac.cn/omix/release/OMIX002290; 11. Mouse uterine tissue dataset (MPSTA): https://db.cngb.org/stomics/mpsta/download/; 12. Mouse embryonic dataset (MOSTA): https://db.cngb.org/stomics/mosta/download/; 13. Injured axolotl telencephalon dataset (ARTISTA): https://db.cngb.org/stomics/artista/download/.

## Code availability

CODA is available at: https://github.com/xiaojierzi/CODA.

## AUTHOR CONTRIBUTIONS

Yecheng Tan: Conceptualization, Methodology, Formal analysis, Visualization, Writing—original draft, Writing—review & editing. Zezhou Wang: Formal analysis, Visualization, Writing—original draft, Writing—review & editing. Ai Wang: Writing— review & editing. Yan Yan: Investigation. Wei Lin: Supervision. Qing Nie: Conceptualization, Methodology, Supervision, Writing—original draft, Writing— review & editing. Jifan Shi: Conceptualization, Methodology, Formal analysis, Visualization, Supervision, Writing—original draft, Writing—review & editing.

## ACKNOWLEDGEMENTS

We thank Liuqian Guo from Dalian University of Technology for her valuable suggestions on drawing the elements for Figure 1(a) and Figure 4(a). This work was supported by National Natural Science Foundation of China [Nos. 12301620, 42450192, 11925103, 82070463], Science and Technology Commission of Shanghai Municipality [No. 21DZ1201402], the AI for Science Foundation of Fudan University [No. FudanX24AI041], and the Shanghai Municipal Data Bureau special fund for urban digital transformation [No. 202401065].

## CONFLICT OF INTEREST

The authors declare no competing interests.

## Notes

### Competing Interest Statement

The authors have declared no competing interest.

