## Supplemental Information for "Integrative cross-sample alignment and spatially differential gene analysis for spatial transcriptomics"

Yecheng Tan[1,2,3], Zezhou Wang[1,4], Ai Wang[5], Yan Yan[5], Wei Lin[1,6,*], Qing Nie[7,*] and Jifan Shi[1,3,6,*]
[1] Research Institute of Intelligent Complex Systems, Fudan University, Shanghai, 200433, China
[2] Institute of Science and Technology for Brain-Inspired Intelligence, Fudan University, Shanghai, 200433, China
[3] Center for Integrative Spatial-Omics Research, Fudan University, Shanghai, 200433, China
[4] Shanghai Center for Mathematical Sciences, Fudan University, Shanghai, 200433, China
[5] Department of Cardiology, Zhongshan Hospital, Fudan University, Shanghai, 200032, China
[6] State Key Laboratory of Medical Neurobiology and MOE Frontiers Center for Brain Science, Fudan University, Shanghai, 200032, China
[7] Department of Mathematics, University of California Irvine, CA, 92697, USA

**1 Supplementary notes**
- **1.1 Datasets and preprocessing.**
  - **1.1.1 Human dorsolateral prefrontal cortex dataset (10X Visium)**
  - **1.1.2 Whole mouse brain cell-type atlas dataset (MERFISH)**
  - **1.1.3 Maize ear primordium dataset (Stereo-seq)**
  - **1.1.4 Mouse coronal brain section dataset (10X Visium)**
  - **1.1.5 Mouse Alzheimer's disease brain dataset (STARmap PLUS)**
  - **1.1.6 Mouse brain spatial translation dataset (RIBOmap)**
  - **1.1.7 Human artery dataset (10X Visium)**
  - **1.1.8 Mouse hippocampus and cortex dataset (CosMx SMI)**
  - **1.1.9 Mouse brain sleep modulation dataset (10X Visium)**
  - **1.1.10 Soybean root nodule dataset (Stereo-seq)**
  - **1.1.11 Mouse uterine tissue dataset (Stereo-seq)**
  - **1.1.12 Mouse embryonic dataset (Stereo-seq)**
  - **1.1.13 Injured axolotl telencephalon dataset**
- **1.2 Identification of the common domain**

**2 Supplementary tables**
- **2.1 Evaluation metrics for DLPFC (Visium) dataset**
- **2.2 Evaluation metrics for Mouse brain (MERFISH) dataset**
- **2.3 Evaluation metrics for developing maize ear (Stereo-seq) dataset**
- **2.4 Spatially consistent genes (SCGs) in the human artery dataset**
- **2.5 Spatially differential genes (SDGs) in the human artery dataset**
- **2.6 Spatially consistent genes (SCGs) in the mouse brain sleep dataset**
- **2.7 Spatially consistent genes (SCGs) in the injured axolotl dataset**

**3 Supplementary figures**

**3.1 Original slices used in the manuscript**

**3.2 Spatial expression of shared SVGs across technologies (before alignment)**

**3.3 Spatial expression of shared SVGs across technologies (after alignment)**

**3.4 Visualization of high cross-correlation marker genes in the AS artery dataset**

**3.5 Visualization of high cross-correlation marker genes in the normal artery dataset**

**3.6 Visualization of spatially differential genes (SDGs) in the artery dataset**

**3.7 Visualization of spatially consistent genes (SCGs) in the mouse brain sleep dataset (Saline)**

**3.8 Visualization of spatially consistent genes (SCGs) in the mouse brain sleep dataset (RHY)**

**3.9 Visualization of spatially consistent genes (SCGs) in the injured axolotl dataset (DPI2 replicate1)**

**3.10 Visualization of spatially consistent genes (SCGs) in the injured axolotl dataset (DPI2 replicate2)**

### Supplementary notes

#### 1.1 Datasets and preprocessing.

We collected nine spatial transcriptomics datasets, covering different species and sequencing technologies, including human dorsolateral prefrontal cortex dataset, whole mouse brain cell-type atlas dataset, maize ear primordium dataset, mouse coronal brain section dataset, mouse Alzheimer's disease brain dataset, mouse brain spatial translation dataset, human artery dataset, mouse hippocampus and cortex dataset and mouse brain sleep modulation dataset.

##### 1.1.1 Human dorsolateral prefrontal cortex dataset (10X Visium)

The human dorsolateral prefrontal cortex (DLPFC) dataset was obtained from KR Maynard et al. [1], and sequenced by 10X Visium. The dataset consists of 12 coronal slices from three independent adult donors without neurological symptoms, with each donor contributing four slices:

- Donor 1: ID151507, ID151508, ID151509, ID151510
- Donor 2: ID151669, ID151670, ID151671, ID151672
- Donor 3: ID151673, ID151674, ID151675, ID151676

Each slice contains approximately 3,000 spatial spots, and the original sequencing captured about 30,000 genes per slice. During initial quality control, mitochondrial genes and low-quality genes were removed, leaving approximately 20,000 genes for further analysis.

In the benchmark section of our manuscript, we selected slices 151673 and 151676 for analysis and performed the following preprocessing steps. First, each dataset was normalized to 10,000 total counts per spot, followed by a log-transformation to normalize gene expression values. Next, we selected the top 3,000 highly variable genes. Then we used ComBat to correct the potential batch effects between slices.

##### 1.1.2 Whole mouse brain cell-type atlas dataset (MERFISH)

The whole mouse brain cell-type atlas dataset is part of the Allen Brain Cell (ABC) Atlas, an open science resource developed by the Allen Institute as part of the Brain Knowledge Platform [2]. This dataset was generated using MERFISH and includes approximately 4 million cells after segmentation and quality control. A 550-gene panel was used to provide high-resolution spatial annotations at multiple hierarchical levels.

In our benchmark, we selected two adjacent brain sections labeled C57BL6J-638850.38 and C57BL6J-638850.40 for analysis, utilizing all 550 genes from the dataset. To account for batch effects between the two sections, we applied ComBat for data integration.

##### 1.1.3 Maize ear primordium dataset (Stereo-seq)

The maize ear primordium dataset was generated using Stereo-seq to construct a high-resolution spatial transcriptomic map of developing maize ear primordium [3]. This dataset includes four sections, with bin and gene counts as follows: the first section contains 8,857 bins and 27,302 detected genes, the second section contains 9,348 bins and 27,302 detected genes, the third section contains 7,139 bins and 22,700 detected genes, and the fourth section contains 8,984 bins and 24,233 detected genes.

In our benchmark, we selected the first and second sections for analysis. We selected the top 3,000 HVGs and used ComBat to correct batch effects between the two slices.

##### 1.1.4 Mouse coronal brain section dataset (10X Visium)

The mouse brain dataset was generated using the 10X Visium platform to spatially profile gene expression across multiple brain regions in adult mice [4]. This dataset includes five coronal sections named ST8059048, ST8059049, ST8059050, ST8059051, and ST8059052. Each section contains transcriptome-wide data across 31,053 genes. The number of spatial bins (spots) per section is as follows: ST8059048 contains 2,987 bins, ST8059049 contains 3,499 bins, ST8059050 contains 3,497 bins, ST8059051 contains 2,409 bins, and ST8059052 contains 2,576 bins.

In our experiments, we used all five sections to evaluate the robustness of spatial alignment and cross-section consistency. We selected all genes and used BBKNN to correct batch effects.

##### 1.1.5 Mouse Alzheimer's disease brain dataset (STARmap PLUS)

The Alzheimer's disease mouse brain dataset was generated using STARmap PLUS [5], an advanced image-based in situ spatial transcriptomics method that enables simultaneous detection of RNA and protein markers at subcellular resolution. The dataset consists of brain sections from TauPS2APP triple transgenic mice, a model exhibiting both amyloid-β (Aβ) plaque and hyperphosphorylated tau (p-tau) pathologies.

We selected two coronal brain sections for analysis, labeled as sample 9494 (8 months) and sample 11346 (13 months), containing 10372 and 9634 cells respectively. Each section was profiled using STARmap PLUS with a curated gene panel of 2,766 genes. These sections encompass cortical and hippocampal regions with prominent Aβ plaques and tau pathology.

In our experiment, we used all genes and used BBKNN to correct batch effects between the two slices.

##### 1.1.6 Mouse brain spatial translation dataset (RIBOmap)

The mouse brain spatial translation dataset was generated using RIBOmap [6], a three-dimensional in situ spatial translatomics method capable of detecting ribosome-bound mRNAs at single-cell and subcellular resolution. We analyzed two coronal hemibrain sections, which together encompass 119,173 cells across major brain regions including cortex, hippocampus, thalamus, hypothalamus, and fiber tracts. Each slice was profiled for 3,034 genes.

In our experiment, we used all genes and used BBKNN to correct batch effects between the two slices.

**1.1.7 Human artery dataset (10X Visium)**

The human artery dataset was generated using 10X Visium spatial transcriptomics to study spatial gene expression patterns in arteries from patients with and without atherosclerosis (AS) [7]. Arterial tissue sections were collected during coronary artery bypass grafting or heart transplantation surgeries, with matched hematoxylin and eosin (H&E) staining used to assess morphological differences. We analyzed two spatial transcriptomics slices: one from normal arteries and the other from an atherosclerotic artery.

In our experiment, we selected the top 3,000 HVGs and used Seurat to correct batch effects between the two slices.

**1.1.8 Mouse hippocampus and cortex dataset (CosMx SMI)**

The mouse brain dataset was generated using the CosMx™ Spatial Molecular Imager (SMI) [8], a single-molecule imaging platform enabling high-resolution spatial transcriptomic profiling in FFPE tissue. The dataset includes two coronal slices: one covering a full hemisphere and the other encompassing the hippocampus and cortex bilaterally.

The coronal hemisphere section contains 48,011 cells, while the hippocampus and cortex section includes 38,987 cells. Each section contains transcriptome-wide data across 950 genes.

In our experiment, we used all genes and used Combat to correct batch effects between the two slices.

**1.1.9 Mouse brain sleep modulation dataset (10X Visium)**

The mouse brain sleep modulation dataset was generated using the 10X Visium Spatial Gene Expression platform to investigate the effects of rhynchophylline (RHY), a natural alkaloid, on sleep-related brain activity and transcriptomic regulation [9]. We selected two coronal brain sections for analysis: GSM6704280 (ZT4_F_Saline) as control and GSM6704281 (ZT4_F_RHY) as the RHY-treated group, both sampled at Zeitgeber Time 4 (ZT4).

Each slice covers the cerebral cortex, hippocampus, and hypothalamic regions associated with sleep regulation, and was profiled using spatial transcriptomics followed by RNA sequencing.

In our experiment, we selected the top 3,000 HVGs and used ComBat to correct batch effects between the two slices.

**1.1.10 Soybean root nodule dataset (Stereo-seq)**

The soybean root nodule dataset was generated using the Stereo-seq spatial transcriptomics platform to capture the spatial organization and transcriptional dynamics of nodules during development [10]. The dataset includes fresh root nodules sampled at two key stages—12 days post-infection (dpi) and 21 dpi—representing early and mature stages of symbiosis.

In our experiments, we selected the top 3,000 HVGs and used ComBat to correct batch effects between the two slices.

###### 1.1.11 Mouse uterine tissue dataset (Stereo-seq)

The mouse uterine dataset was generated using the Stereo-seq platform to construct a high-resolution spatiotemporal transcriptomic atlas of mouse placentation (MPSTA) [11]. This dataset spans developmental stages from embryonic day (E) 7.5 to E14.5. In our benchmark, we selected two sagittal uterine sections at E8.5, named E8.5_S1.MPSTA and E8.5_S2.MPSTA, both capturing the fetal-maternal interface during early placental development.

In our experiment, we selected the top 3,000 HVGs and used BBKNN to correct batch effects between the two slices.

###### 1.1.12 Mouse embryonic dataset (Stereo-seq)

The mouse embryonic dataset was generated using Stereo-seq, a high-resolution spatial transcriptomics platform based on DNA nanoball (DNB) patterned arrays [12]. The dataset is part of the MOSTA (Mouse Organogenesis Spatiotemporal Transcriptomic Atlas) project and covers key developmental stages from E9.5 to E16.5.

In our benchmark, we selected two sagittal embryo sections at embryonic day 15.5 (E15.5_E1S3 and E15.5_E1S4), both representing mid-gestation stages with extensive tissue patterning.

In our experiment, we selected the top 1,000 HVGs and used BBKNN to correct batch effects between the two slices.

###### 1.1.13 Injured axolotl telencephalon dataset (Stereo-seq)

The axolotl telencephalon dataset was generated using Stereo-seq, a high-resolution spatial transcriptomics technology capable of single-cell resolution mapping [13]. This dataset captures the regenerative process of the axolotl brain following injury to the dorsal pallium region of the left telencephalic hemisphere. Tissue sections were collected at seven time points post-injury—2, 5, 10, 15, 20, 30, and 60 days (DPI).

In our experiment, we selected coronal sections at 2 DPI to investigate early injury response and mid-stage regeneration, chose the top 3,000 HVGs and used BBKNN to correct batch effects between the two slices.

##### 1.2 Identification of the common domain

After establishing correspondences between feature points $\mathcal{P}_{\mathrm{S}} = \{I_S(x_i^S, y_i^S)\}_{i=1}^{l}$ and $\mathcal{P}_{\mathrm{T}} = \{I_T(x_i^T, y_i^T)\}_{i=1}^{l}$ in the source and target slices using the transformer-based attention module, CODA provides two alternative strategies to define the common domain, based on the spatial location of matched keypoints. These options offer trade-offs between simplicity and geometric robustness, and their detailed implementation is as follows.

**Axis-Aligned Bounding Box Strategy**. This default strategy selects the overlapping rectangular region that minimally encloses all matched keypoints. The common domain is then defined based on the row and column coordinates of the feature pixels:

$$R_S = \{(x, y) | x^S_{\min} \le x \le x^S_{max}, y^S_{min} \le y \le y^S_{max}\}$$

$$R_T = \{(x, y) | x^T_{\min} \le x \le x^T_{max}, y^T_{min} \le y \le y^T_{max}\}$$

where $x^S_{\min}$, $x^T_{\min}$, $y^S_{min}$ and $y^T_{min}$ represent the minimum $x$ and $y$ coordinates of the corresponding pixels in the source and target slices, and $x^S_{max}$, $x^T_{max}$, $y^S_{max}$ and $y^T_{max}$ represent the maximum $x$ and $y$ coordinates of the corresponding pixels in the source and target slices, respectively.

This method is simple and fast, but may include redundant boundary regions or exclude irregular overlaps.

**Convex Hull Intersection Strategy**. To improve geometric precision and reduce the influence of outliers, CODA also supports a convex hull–based common domain. We compute the convex hulls of matched keypoints in both slices:

$$R_S = \mathrm{Conv}(\mathcal{P}_\mathrm{S}), \qquad R_T = \mathrm{Conv}(\mathcal{P}_\mathrm{T})$$

where denotes the convex hull operation. The common domain is then defined as the geometric intersection:

This region tends to be more biologically meaningful and spatially compact.

**Cell Selection**. Once is identified using either strategy, CODA selects only those cells whose spatial coordinates fall within this region. Let and denote all spatial coordinates in the source and target slices. Then:

$$\mathcal{X}'_\mathrm{S} = \{x^S_i \in \mathcal{X}_\mathrm{S} | x^S_i \in R_S\}, \qquad \mathcal{X}'_\mathrm{T} = \{x^T_j \in \mathcal{X}_\mathrm{T} | x^T_j \in R_T\}$$

#### Supplementary tables

##### 2.1 Evaluation metrics for DLPFC (Visium) dataset

| | Before alignment | CODA (CPU) | PASTE (GPU) | SANTO (CPU) | STalign (CPU) | SA-GEARS (CPU) |
|---|---|---|---|---|---|---|
| ECS | 0.30 | 0.37 | 0.32 | 0.34 | 0.30 | 0.37 |
| SCI | 0.18 | 0.84 | 0.35 | 0.37 | 0.18 | 0.86 |
| RRS | 0.00 | 0.98 | 0.73 | 0.76 | 0.00 | 0.99 |
| Running time | | 58.38 s | 113.15 s | 210.10 s | 213.72 s | 287.41 s |
| Memory use | | 947.89 MB (RAM) | 1104.02 MB (RAM) + 925.42 MB (VRAM) | 823.66 MB (RAM) | 72.62 MB (RAM) | 1097.19 MB (RAM) |

**Table 1. Evaluation metrics for DLPFC (Visium) dataset.** Comparison of global spatial alignment performance across five methods on the DLPFC dataset: CODA, PASTE, SANTO, STalign, and ST-GEARS.

##### 2.2 Evaluation metrics for Mouse brain (MERFISH) dataset

| | Before alignment | CODA (CPU) | PASTE (GPU) | SANTO (CPU) | STalign (CPU) | SA-GEARS (CPU) |
|---|---|---|---|---|---|---|
| ECS | 0.32 | 0.43 | 0.36 | 0.30 | 0.32 | 0.41 |
| SCI | 0.18 | 0.29 | 0.20 | 0.14 | 0.17 | 0.27 |
| RRS | 0.00 | 0.97 | 0.84 | 0.99 | 0.00 | 0.99 |
| Running time | | 509.95 s | 949.96 s (On sampled 10,000 cells) | 1224.50 s (On sampled 10,000 cells) | 239.35s | 4930.41 s (On sampled 10,000 cells) |
| Memory use | | 1245.51 MB (RAM) | 1370.54 MB (RAM) + 6665.08 MB (VRAM) | 6772.13 MB (RAM) | 58.20 MB (RAM) | 1265.90 MB (RAM) |

**Table 2. Evaluation metrics for Mouse brain (MERFISH) dataset.** Comparison of global spatial alignment performance across five methods on the Mouse brain (MERFISH) dataset: CODA, PASTE, SANTO, STalign, and ST-GEARS.

##### 2.3 Evaluation metrics for developing maize ear (Stereo-seq) dataset

| | Before | CODA | PASTE | SANTO | STalign | SA-GEARS |
|---|---|---|---|---|---|---|

| | alignment | (CPU) | (GPU) | (CPU) | (CPU) | (CPU) |
|---|---|---|---|---|---|---|
| ECS | 0.55 | 0.70 | 0.59 | 0.70 | 0.55 | 0.58 |
| SCI | 0.08 | 0.38 | 0.13 | 0.40 | 0.08 | 0.10 |
| Running time | | 99.08 s | 1023.71 s | 919.07 s | 213.98 s | 3767.31 s |
| Memory use | | 599.03 MB (RAM) | 1504.10 MB (RAM) + 5680.21 MB (VRAM) | 713.33 MB (RAM) | 67.92 MB (RAM) | 1244.89 MB (RAM) |

**Table 3. Evaluation metrics for developing maize ear (Stereo-seq) dataset.** Comparison of global spatial alignment performance across five methods on the developing maize ear (Stereo-seq) dataset: CODA, PASTE, SANTO, STalign, and ST-GEARS.

#### 2.4 Spatially consistent genes (SCGs) in the human artery dataset (top 2% ranked by spatial cross-correlation)

| | | | | | | | | | | |
|---|---|---|---|---|---|---|---|---|---|---|
| MYL9 | TAGLN | ACTA2 | APOD | CFD | FN1 | TPM2 | FLNA | MYH11 | G0S2 | MYL9 |
| DCN | C7 | LINC00632 | MGST1 | FABP4 | AEBP1 | SERPINF1 | DSTN | AL161421.1 | PLIN4 | DCN |
| GSN | BGN | CXCL14 | CNN1 | HSPB1 | C3 | PLIN1 | FBLN1 | RPL37 | ACTG2 | GSN |
| GPD1 | SELENOP | HTRA3 | CD36 | APOE | MGP | IGFBP4 | SCARA5 | S100B | MCAM | GPD1 |
| PLP1 | PLN | PPP1R14A | HLA-B | SCGB3A1 | MUSTN1 | PODN | TGM2 | ADAMTS13 | SORBS2 | PLP1 |
| HEMGN | HOTAIRM1 | FBLN2 | CCDC3 | NAV3 | TAF5 | ACTB | RPL17 | LIPE | DPT | HEMGN |

**Table 4. Spatially consistent genes (SCGs) in the human artery dataset.** List of the top 2% genes ranked by spatial cross-correlation score across spatially aligned slices in the human artery dataset. These genes exhibit highly consistent spatial expression patterns across samples and are potential markers of preserved spatial organization. Many of the top-ranked SCGs, such as MYL9, TAGLN, and ACTA2, are known to be associated with vascular smooth muscle cells and arterial structure.

#### 2.5 Spatially differential genes (SDGs) in the human artery dataset (bottom 2% ranked by spatial cross-correlation)

| | | | | | | | | | | |
|---|---|---|---|---|---|---|---|---|---|---|
| TYMP | ZDBF2 | POLR2F | PASK | HCLS1 | SGSM3 | ACE | SLC25A16 | EFL1 | SCN3A | TYMP |
| TTC39C | IGHG2 | RASGRP3 | SLC25A28 | LY96 | TRIM7 | KCNK15 | KRI1 | ARHGAP45 | PARVB | TTC39C |
| N4BP1 | WRAP73 | AC010969.2 | QKI | WDR36 | ENDOV | KLF11 | FLVCR1-DT | TFPT | IGLC1 | N4BP1 |
| DPM2 | PCDH20 | CD180 | ATP5MC1 | REX1BD | SMARCD3 | CTSD | MAP2K5 | LACC1 | JRK | DPM2 |
| CSGALNACT1 | CTSF | ALS2CL | BPHL | RMND1 | AL135910.1 | ARHGEF26-AS1 | MTSS1 | FAAP100 | KIFC3 | CSGALNACT1 |
| SLAMF7 | AP2A2 | CRYZ | DCP1A | ARHGA | JCHAIN | ARHGEF7 | RARS2 | CCL2 | MKI67 | SLAMF7 |

| | | | | |
|---|---|---|---|---|
| | | | | P4 |

**Table 5. Spatially differential genes (SDGs) in the human artery dataset.** List of the bottom 2% genes ranked by spatial cross-correlation score across spatially aligned slices in the human artery dataset. These genes exhibit condition-specific or spatially divergent expression patterns across samples, indicating potential spatial heterogeneity or localized regulatory shifts. Examples include immune-related (CCL2, SLAMF7), metabolic (TYMP), and signaling genes (MAP2K5, QKI), which may reflect regional or pathological differences in arterial tissue.

#### 2.6 Spatially consistent genes (SCGs) in the mouse brain sleep dataset (top 2% ranked by spatial cross-correlation)

| | | | | | | | | | | |
|---|---|---|---|---|---|---|---|---|---|---|
| Itpka | Baiap3 | AW551984 | Hcrt | Tcf7l2 | Nrgn | Ddn | Slc17a7 | Ngb | Ctxn1 | Itpka |
| Baiap2 | Gpx3 | Slc30a3 | Lamp5 | Gm11549 | Gal | Rprml | Vxn | Gpr151 | Prkcd | Baiap2 |
| Pmch | Hpcal4 | Dlk1 | Igfbp6 | Nrn1 | Psd | Ucn3 | Calb2 | Ngef | Stx1a | Pmch |
| Penk | Arhgap36 | Tbr1 | Shox2 | Gda | Slc17a6 | Snca | Ntng1 | Hap1 | Agt | Penk |
| Mef2c | Zcchc12 | Hpca | Zic1 | Camk2a | Fxyd6 | C1ql2 | Kcnip2 | Nts | Enc1 | Mef2c |
| Cartpt | Sst | Ramp3 | Tnnt1 | Patj | 2010300C02Rik | Rprm | Atp2b4 | Tac1 | Ccn3 | Cartpt |

**Table 6. Spatially consistent genes (SCGs) in the mouse brain sleep dataset.** Top 2% of genes ranked by spatial cross-correlation score in the mouse brain sleep modulation dataset. These genes exhibit highly conserved spatial expression patterns across aligned conditions (e.g., saline vs. RHY treatment), suggesting their spatial organization is preserved during sleep-related modulation. Notable genes include Hcrt and Pmch (key neuropeptides in sleep regulation), Nrgn, Tcf7l2, Slc17a7 (neuronal markers), and Sst, Cartpt, Tac1 (involved in neurotransmission and circadian processes).

#### 2.7 Spatially consistent genes (SCGs) in the injured axolotl dataset (top 2% ranked by spatial cross-correlation)

| | | | | | | | | | | |
|---|---|---|---|---|---|---|---|---|---|---|
| TRH | AMEX60DD029717 | GFAP | PENK | GAD1 | ATF3 | nan | SNCA | COX1 | MAG | TRH |
| DR999_PMT13074[nr] | LOC101397869[nr] | NTS | ZIC1 | GLUL.L[nr] | SCGN | CD9 | NPTX2 | CPEOMESODERMIN[nr] | LOC108785612[nr] | DR999_PMT13074[nr] |
| LOC112114046[nr] | RGCC[nr] | LFNG | LOC115086130[nr] | AMEX60DD017847 | AMEX60DD054004 | ST6GALNAC5 | NMB | LOC115479139[nr] | SST | LOC112114046[nr] |
| SFRP1 | ATP1A2[hs] | TF-A[nr] | RGR | PCP4L1[nr] | CPLX3 | NPTX1 | NPPC4[nr] | ADCYAP1 | NELL2 | SFRP1 |
| LOC115460661[nr] | SLC4A4 | PTN[nr] | GLUL-LIKE.1[nr] | LOC113746826[nr] | GLUD1 | HMOX1 | AMEX60DD035705 | KIAA0513 | S10AB[nr] | LOC115460661[nr] |
| EDNRB | FABP7[nr] | CXCL14.L[nr] | H355_001130[nr] | SYN[nr] | VIM | GJA1 | SNAP25 | SLC26A5 | AB205_0156120[nr] | EDNRB |

**Table 7. Spatially consistent genes (SCGs) in the injured axolotl dataset.** Top 2%

of genes ranked by spatial cross-correlation score in the regenerating axolotl telencephalon dataset at 2 days post injury (2 DPI). These genes show conserved spatial expression across sections, indicating involvement in shared regenerative programs. Notable genes include GFAP, VIM, and FABP7 (markers of reactive glia), GAD1, SNCA, SST (neuronal lineage markers), as well as TRH, EDNRB, SFRP1, RGCC, and PTN, which have been previously implicated in neuroendocrine signaling, injury response, and neural development.

### Supplementary figures

#### 3.1 Original slices used in the manuscript

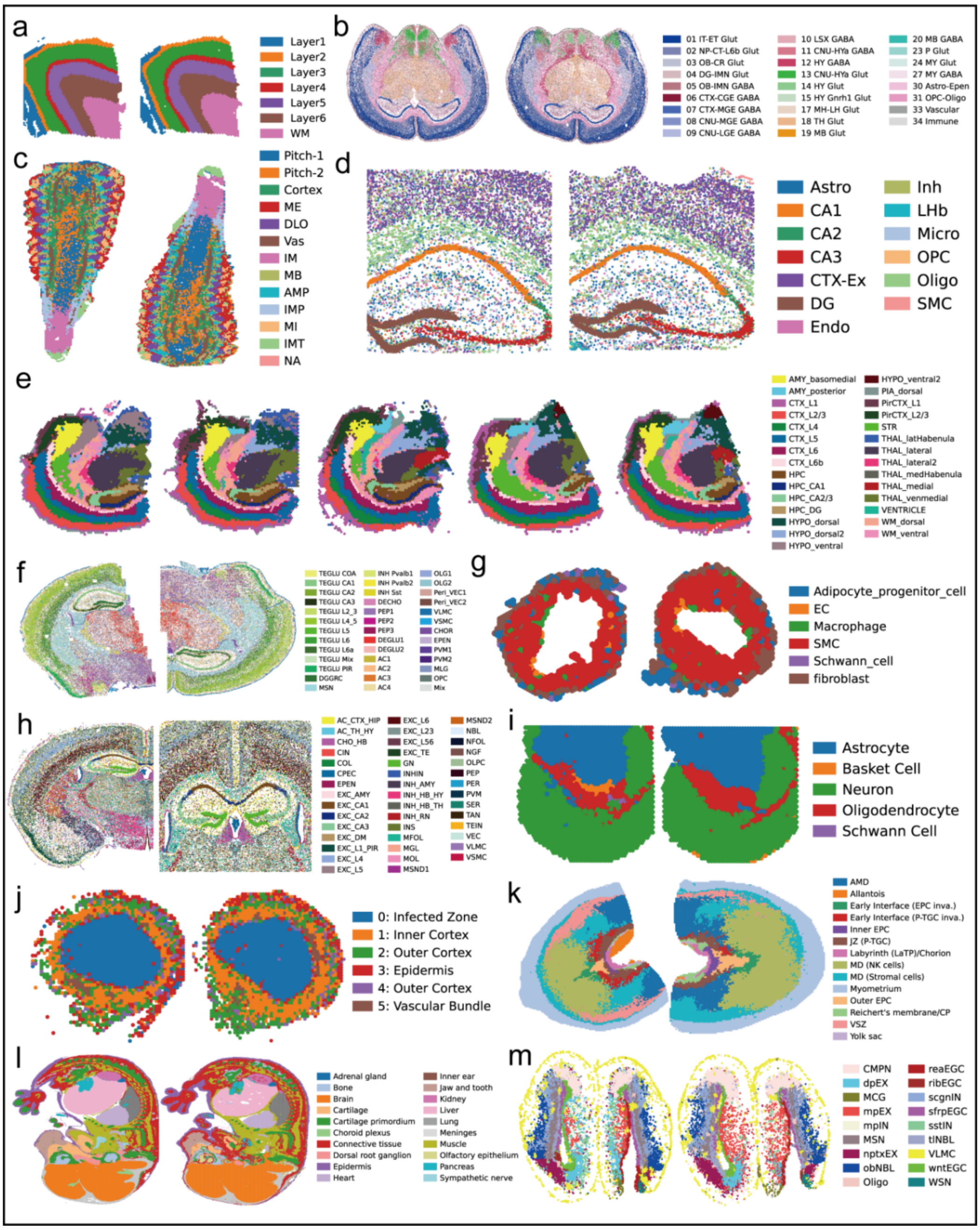

**Figure 1. Slices from all 13 spatial transcriptomics datasets used in this study, spanning diverse species, tissues, platforms, and biological contexts.** (a) human dorsolateral prefrontal cortex (10X Visium), (b) mouse brain (MERFISH), (c) developing maize ear (Stereo-seq), (d) mouse Alzheimer's brain (STARmap PLUS), (e) mouse brain (10X Visium), (f) mouse brain spatial translation dataset (RIBOmap), (g) human artery (10X Visium), (h) mouse brain (CosMx SMI), (i) mouse brain under sleep modulation (10X Visium), (j) soybean root nodule, (k) mouse uterine tissue (MPSTA), (l) mouse embryonic brain (MOSTA), (m) regenerating axolotl telencephalon (ARTISTA).

##### 3.2 Spatial expression of shared SVGs across technologies (before alignment)

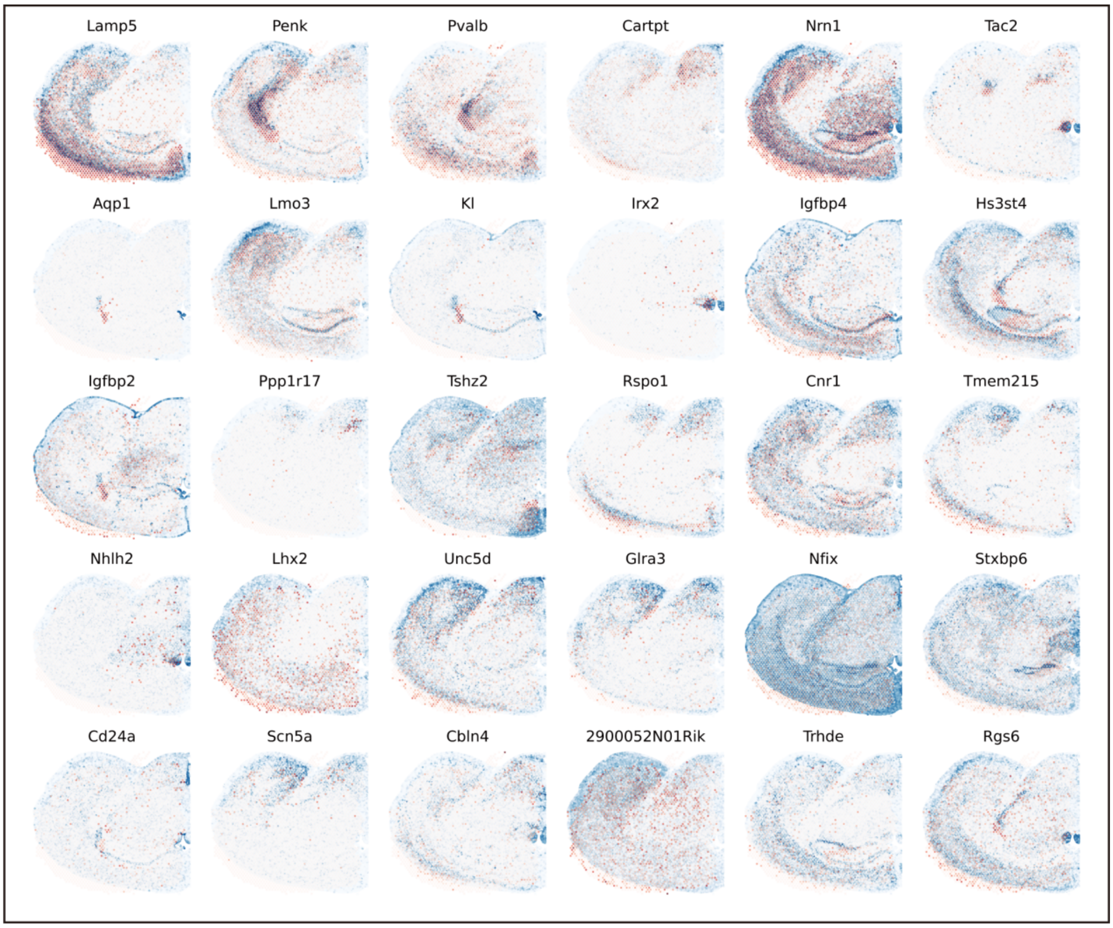

**Figure 2. Spatial expression patterns of selected shared SVGs in slices from different spatial transcriptomics platforms prior to alignment.**

##### 3.3 Spatial expression of shared SVGs across technologies (after alignment)

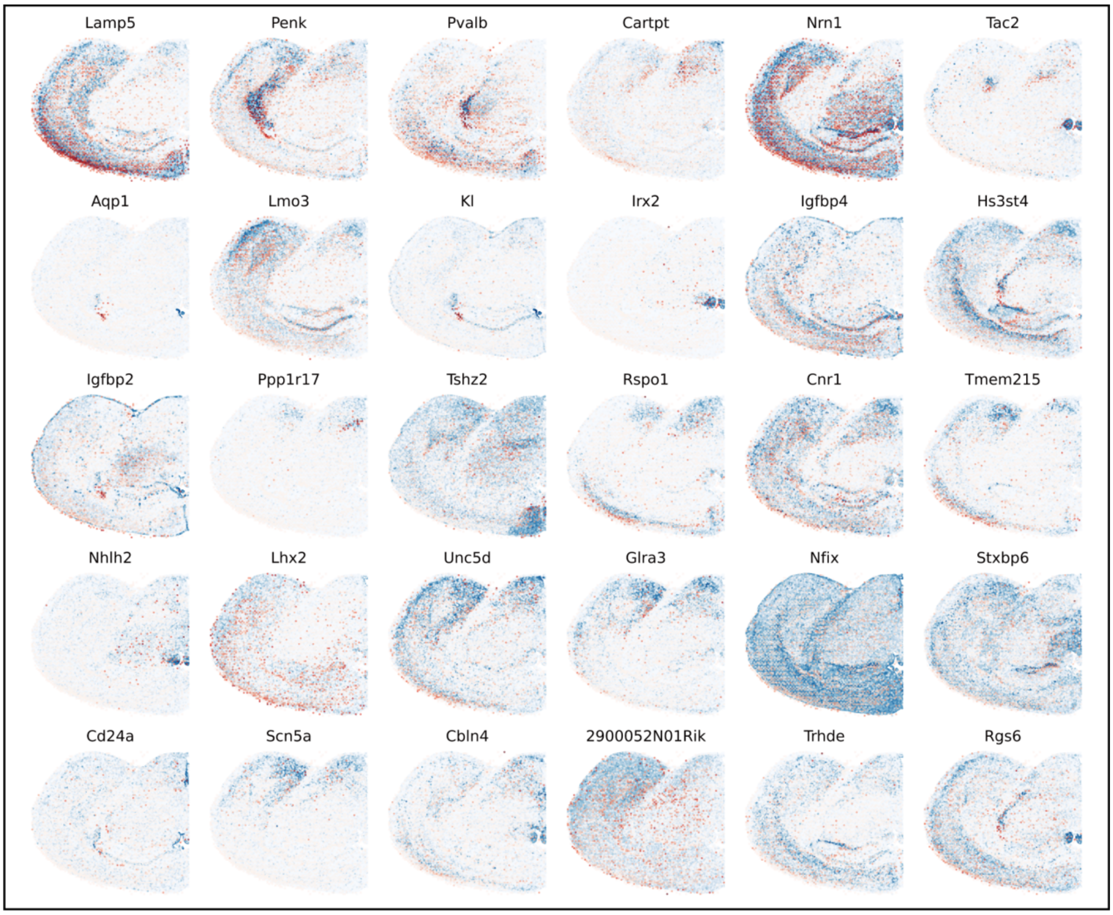

**Figure 3. Spatial expression patterns of selected shared SVGs in slices from different spatial transcriptomics platforms after alignment.**

##### 3.4 Visualization of high cross-correlation marker genes in the AS artery dataset

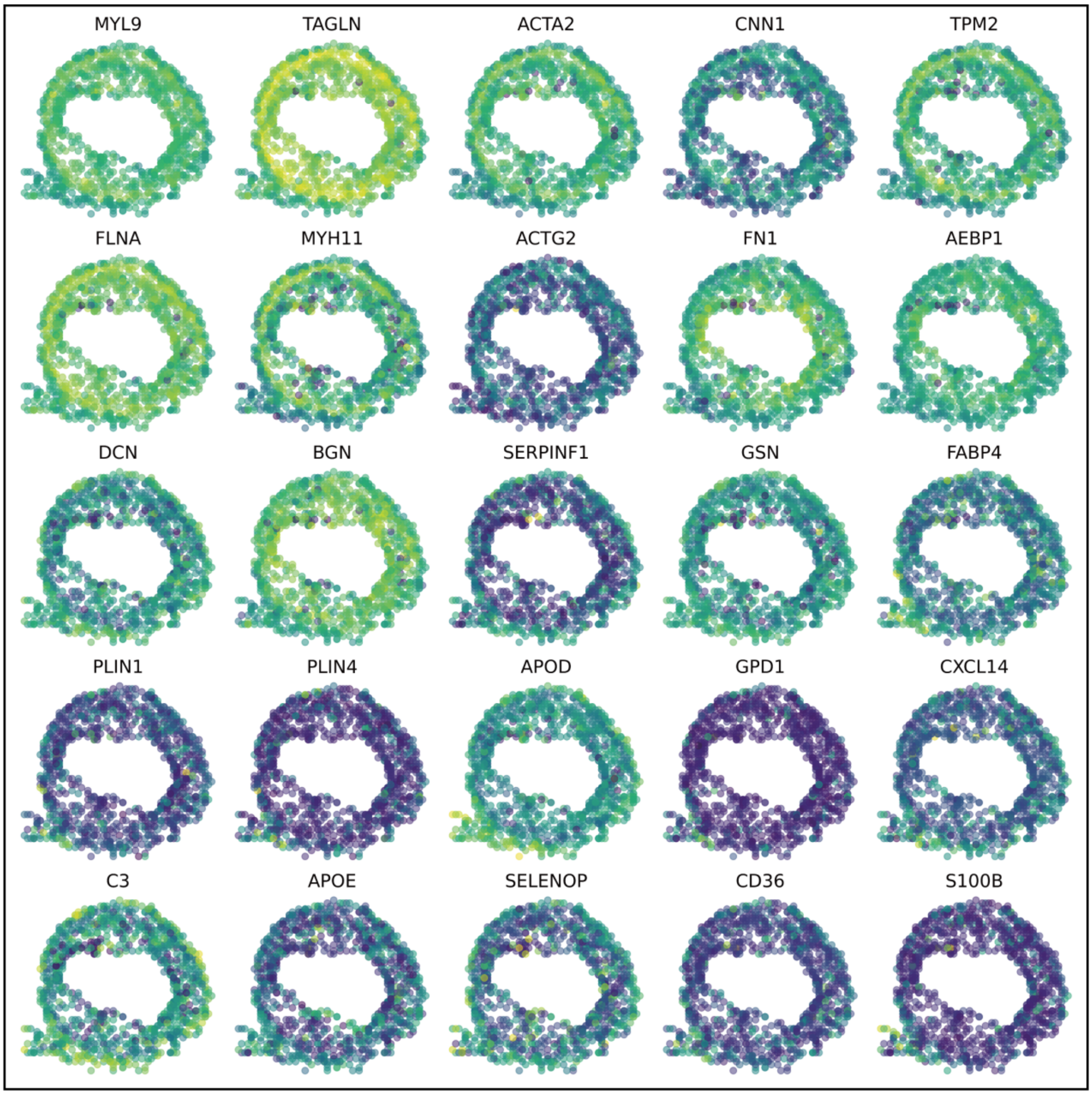

**Figure 4. Visualization of high cross-correlation marker genes in the AS artery dataset.** Spatial expression patterns of selected marker genes with top-ranked spatial cross-correlation scores in the atherosclerotic artery dataset. These genes, including MYL9, TAGLN, ACTA2, and CNN1, reflect conserved smooth muscle cell and extracellular matrix‑related programs associated with vascular pathology. See also Figure 5 for the corresponding expression in normal artery tissue.

##### 3.5 Visualization of high cross-correlation marker genes in the normal artery dataset

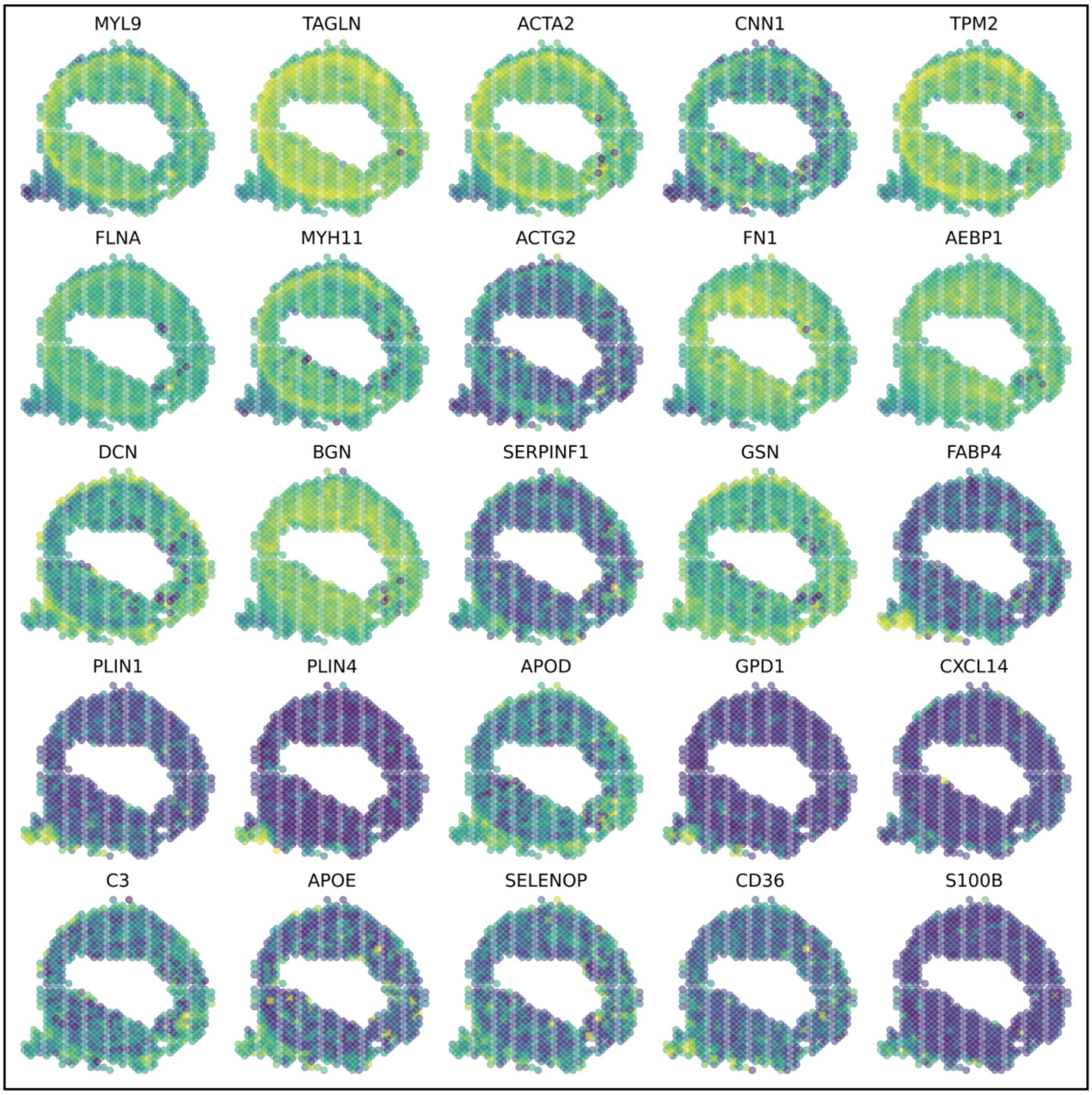

**Figure 5. Visualization of high cross-correlation marker genes in the normal artery dataset.** Spatial expression patterns of the same marker genes (MYL9, TAGLN, ACTA2, and CNN1) in the normal artery dataset. Compared to the atherosclerotic artery (Figure 4), these genes show distinct localization and reduced spatial cross-correlation, suggesting condition-specific remodeling of vascular architecture.

##### 3.6 Visualization of spatially differential genes (SDGs) in the artery dataset

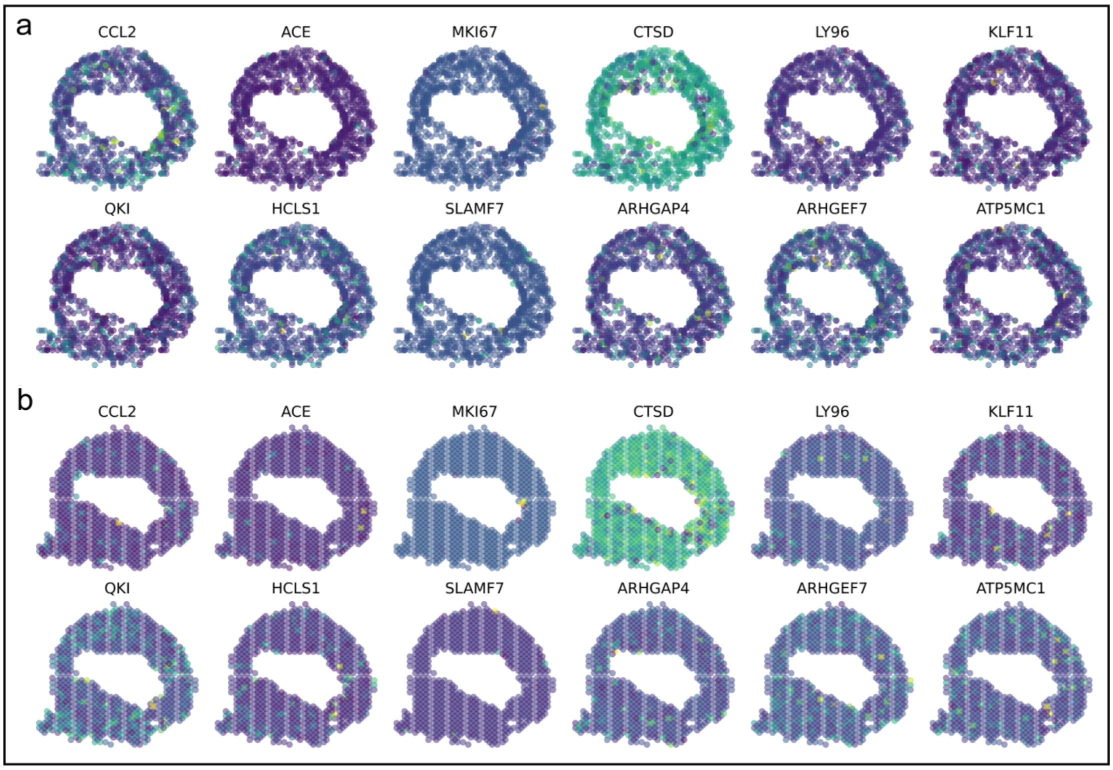

**Figure 6. Visualization of spatially differential genes (SDGs) in the artery dataset.** (a) Spatial expression patterns of representative SDGs in the atherosclerotic artery (CAD). (b) Corresponding spatial expression of the same genes in the normal artery. The selected genes exhibit low spatial cross-correlation between conditions, highlighting spatially heterogeneous expression patterns associated with vascular pathology. These spatially differential genes (SDGs) illustrate CODA's ability to detect condition-specific spatial gene expression differences.

##### 3.7 Visualization of spatially consistent genes (SCGs) in the mouse brain sleep dataset (Saline)

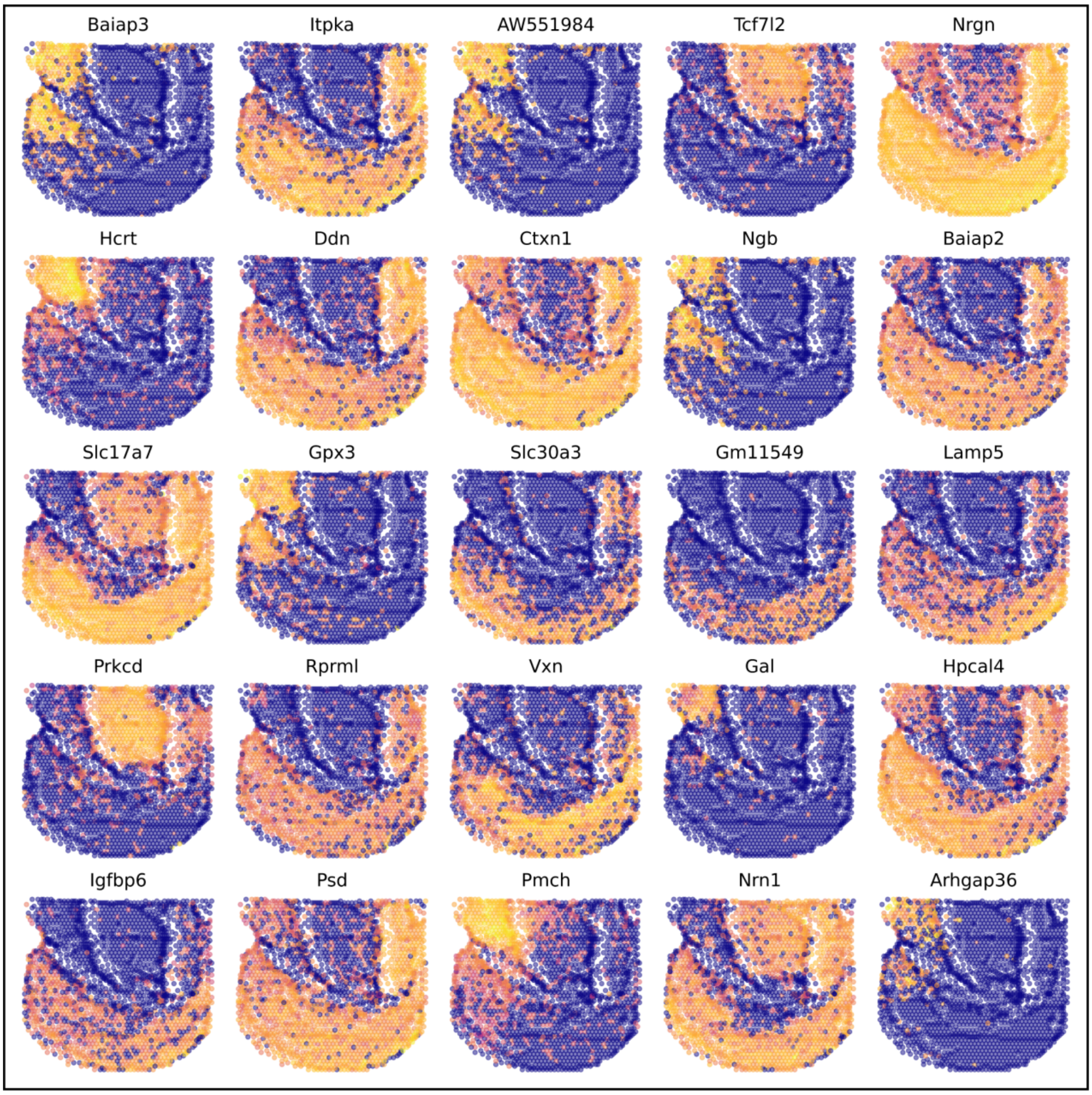

**Figure 7. Visualization of spatially consistent genes (SCGs) in the mouse brain sleep dataset (Saline).** Spatial expression patterns of selected SCGs in the Saline-treated mouse brain. These genes exhibit high spatial cross-correlation and consistent spatial organization across slices.

##### 3.8 Visualization of spatially consistent genes (SCGs) in the mouse brain sleep dataset (RHY)

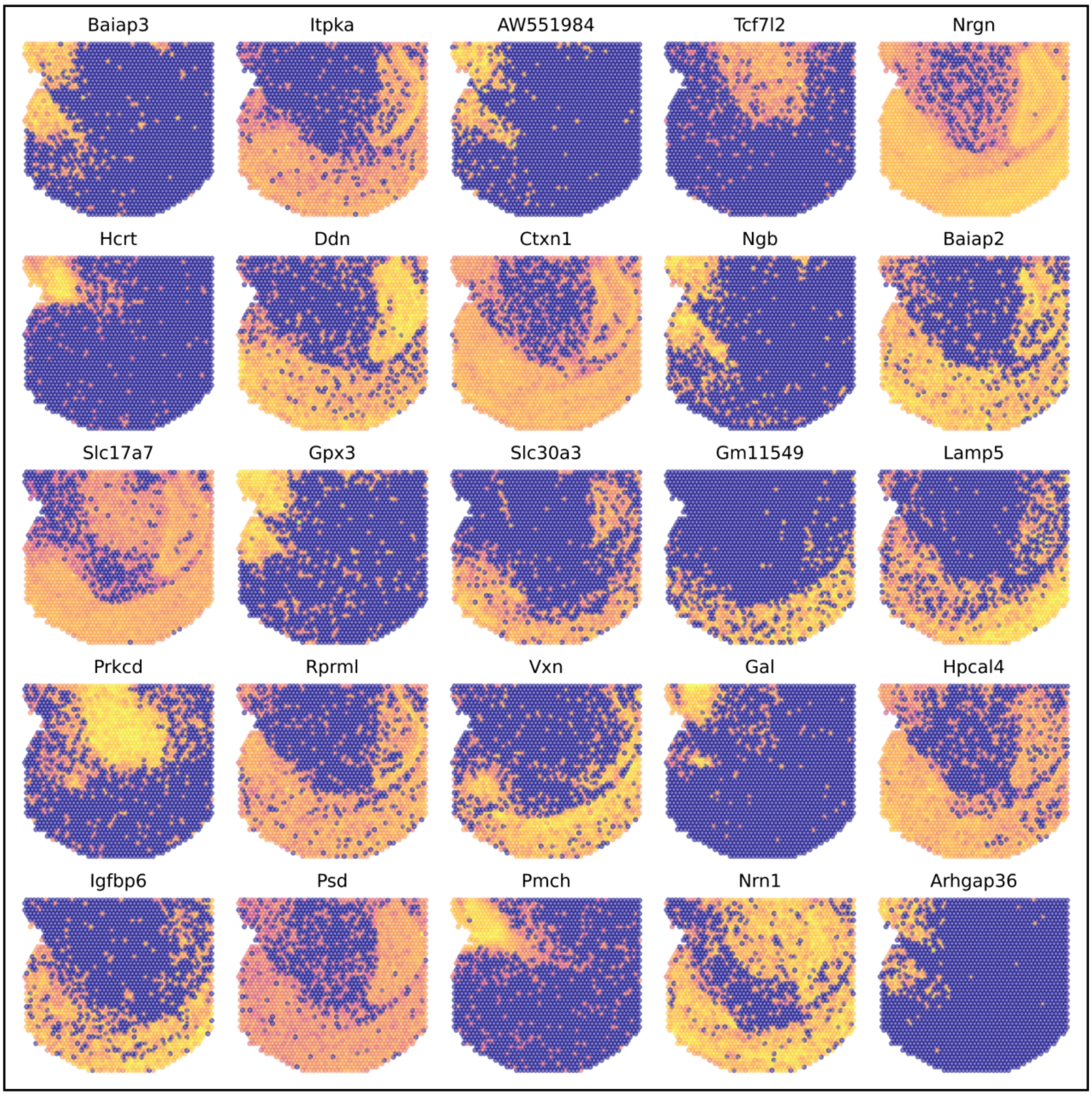

**Figure 8. Visualization of spatially consistent genes (SCGs) in the mouse brain sleep dataset (RHY).** Spatial expression of the same SCGs shown in Figure 7, now in the RHY-treated mouse brain.

##### 3.9 Visualization of spatially consistent genes (SCGs) in the injured axolotl dataset

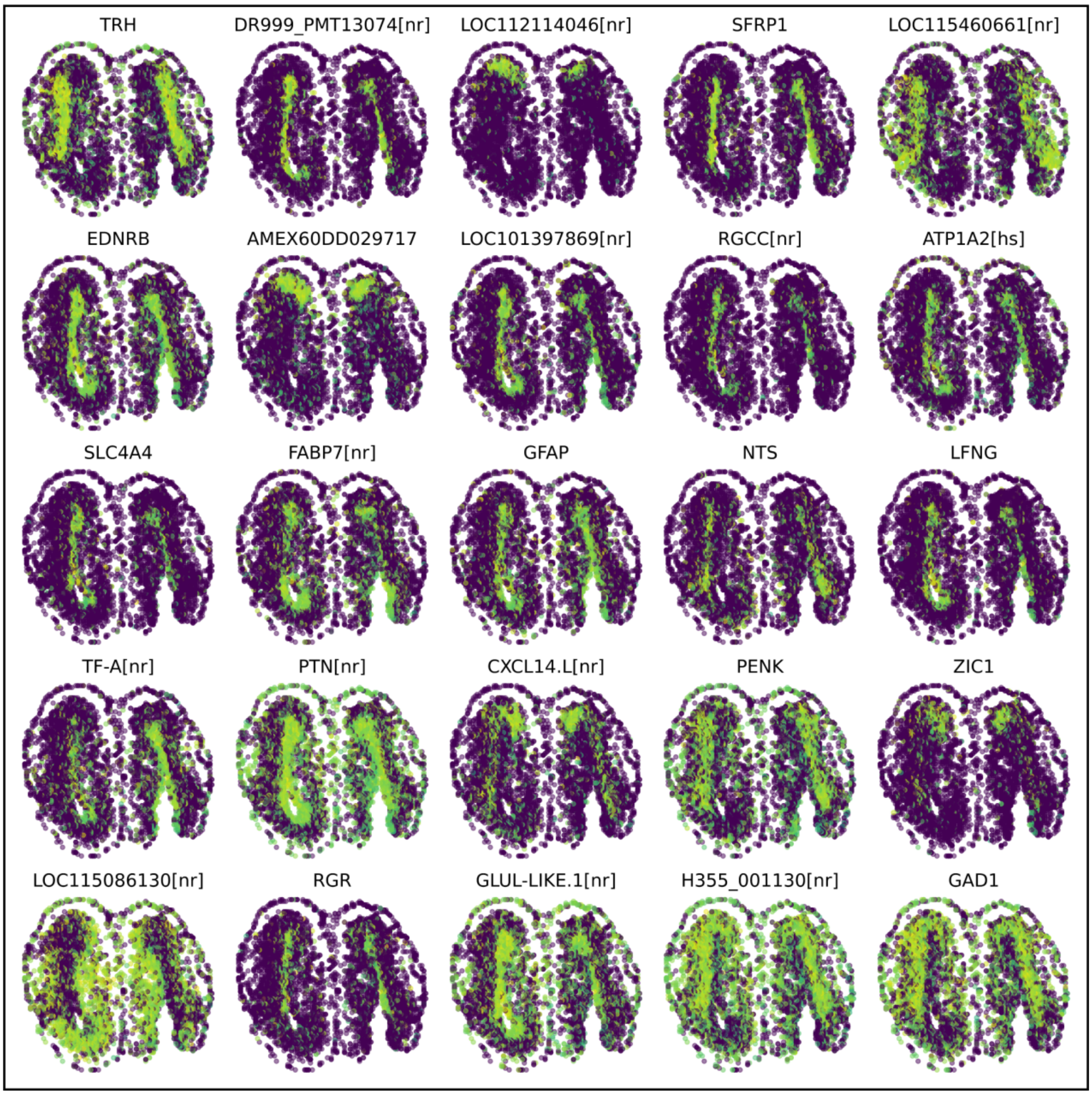

**Figure 9. Visualization of spatially consistent genes (SCGs) in the injured axolotl dataset (2DPI replicate1).** Spatial expression patterns of selected SCGs in the first slice of the injured axolotl telencephalon. These genes, such as TRH, EDNRB, and PTN, exhibit high cross-sample spatial correlation, reflecting conserved spatial programs involved in neuroregeneration and injury response.

##### 3.10 Visualization of spatially consistent genes (SCGs) in the injured axolotl dataset

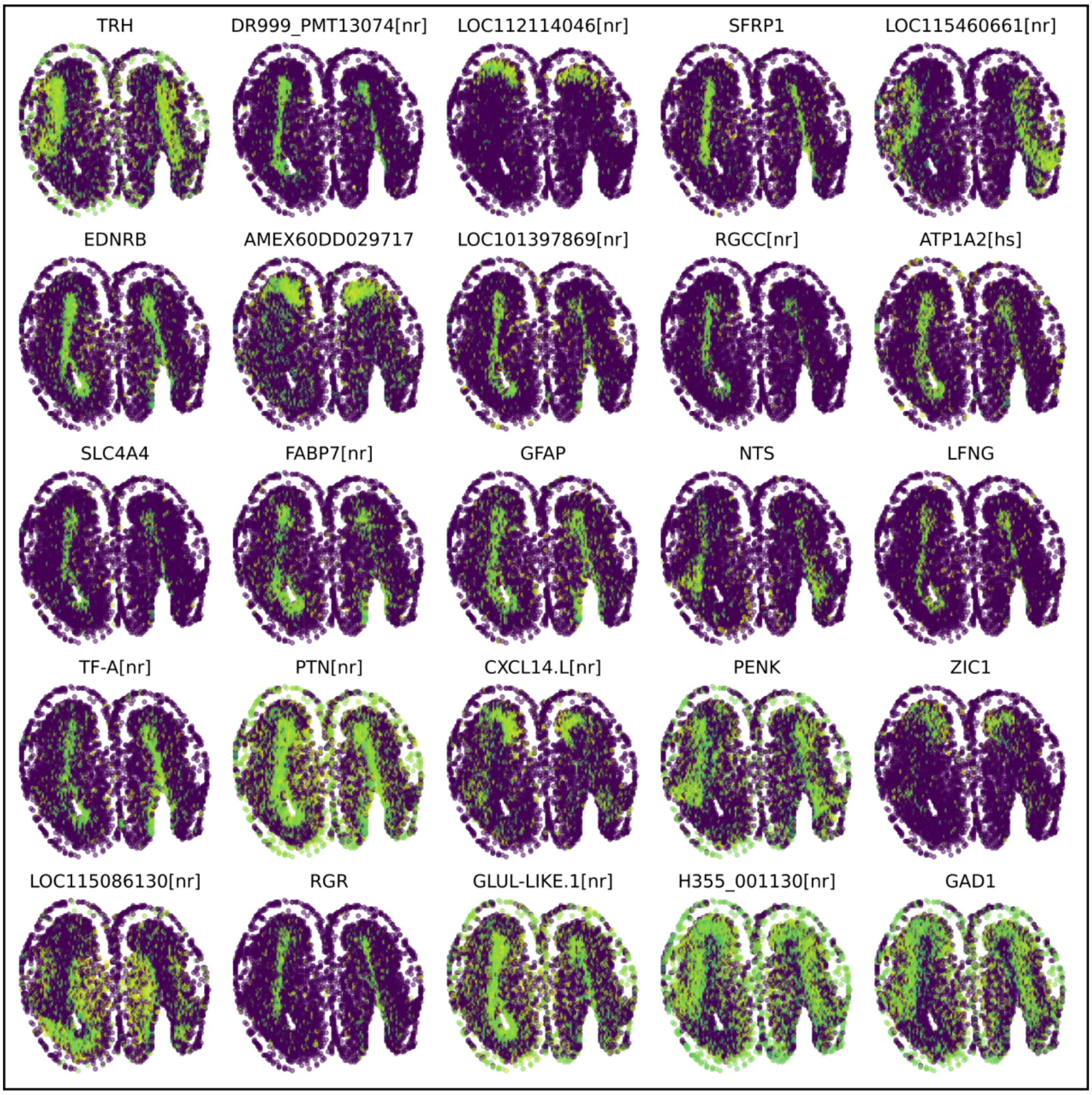

**Figure 10. Visualization of spatially consistent genes (SCGs) in the injured axolotl dataset (2DPI replicate2).** Spatial expression patterns of the same SCGs shown in Figure 3.9, now in the second slice.
